# An axon-intrinsic loop restricts nerve regeneration through axonal protein synthesis

**DOI:** 10.1101/2025.10.28.685020

**Authors:** Courtney N. Buchanan, Jinyoung Lee, Samaneh Matoo, Marie-Claire Honoree, Molly Conway, Lillian F. Thompson, Ansley McKay, Irene Dalla Costa, Moira Lopez De Leon, Elizabeth Thames, Nora Perrone-Bizzozero, Ashley L. Kalinski, Lauren S. Vaughn, Jeffery L. Twiss

## Abstract

Injured axons synthesize the RNA Binding Protein KHSRP that promotes mRNA decay and slows nerve regeneration. Axotomy-induced increase in axoplasmic Ca^2+^ activates axonal *Khsrp* translation, and while Ca^2+^ returns to pre-injury levels within 16 hours post-axotomy, axonal KHSRP remains elevated. Alternating translation of *Reg3a* and *Khsrp* sustains axonal KHSRP levels in regenerating axons. Nerve injury activates *Reg3a* expression, resulting in increased REG3A synthesis and secretion from axons. REG3A stimulates ER Ca^2+^ release to activate PERK, increase eIF2*α* phosphorylation, and increase *Khsrp* translation. Axoplasmic Ca^2+^ slowly oscillates in growth cones and Reg3A depletion attenuates growth cone Ca^2+^ oscillations, decreases KHSRP synthesis, reduces the axon’s retractive events, and accelerates peripheral nerve regeneration. Thus, REG3A to KHSRP signaling provides an axon-intrinsic loop that decelerates axon growth through localized mRNA translation.

## MAIN TEXT

Though injured axons in the peripheral nervous system (PNS) can spontaneously regenerate, the rate of axon growth is quite slow at 1-3 mm per day (*1, 2*). Regeneration over distances of 5 cm or greater is often unsuccessful since the environment within the distal severed nerve loses the ability to support regeneration and target tissues are less receptive for reinnervation (*3*). Accelerating rates of regeneration would allow axons to reach nerve segments well beyond the injury site and innervate target tissues before loss of support and tissues’ receptiveness for reinnervation occur. We recently showed that deletion of the murine KH-type splicing regulatory factor (KHSRP) accelerates nerve regeneration (*4*). KHSRP is a multifunctional RNA binding protein that contributes to RNA splicing, mRNA transport, and mRNA decay (*5*). KHSRP slows axon growth by promoting decay of axonal mRNAs with AU-rich elements (ARE) in their 3’ untranslated regions (UTR). *Khsrp* mRNA translation is acutely activated in injured axons through a Ca^2+^→PERK→eIF2*α*^PS51^ signaling cascade (*4*). However, axonal translation switches from mRNAs that are efficiently translated with elevated axoplasmic Ca^2+^, like *Khsrp* and *Kpnb1*, to mRNAs that are more efficiently translated at lower axoplasmic Ca^2+^, like *Csnk2a1* and *Nrn1*, over 16 h after axotomy (*4, 6*). Despite that switch in translation, axonal KHSRP levels remain elevated in peripheral nerve axons for weeks after axotomy. How are axonal KHSRP levels sustained if its mRNA’s translation requires elevated axoplasmic Ca^2+^? Here, we show that axonal synthesis of the lectin-like protein Regenerating Family Member 3*α* protein (REG3A; also called Pancreatitis-associated protein 2 [PAP2] and Lithostathine 3) (*7*) activates translation of axonal *Khsrp* mRNA in regenerating axons to slow PNS nerve regeneration.

## RESULTS

### REG3A protein slows axon growth

Axonal *Reg3A* mRNA levels increase in regenerating peripheral nerve and ascending spinal cord axons after axotomy (*8*). Consistent with previous findings (*8*), *Reg3a* mRNA increases in sciatic nerve axons at 7 d following crush injury (Figure 1A). Immunoblots of sciatic nerve lysates further show increased REG3A protein over 7-28 d after nerve crush injury (Figure 1B). We used shRNA depletion to determine if the axonal *Reg3A* mRNA increase contributes to axon growth. Cultured adult mouse dorsal root ganglia (DRG) neurons transduced with AAV9-shRNA targeting *Reg3a* (shReg3a) show increased axon growth compared to those transduced with non-targeting AAV9-shRNA (shCntl; Figure 1C,D). Reverse transcriptase coupled droplet digital PCR (RTddPCR) and immunofluorescence confirmed that both *Reg3a* mRNA and REG3A protein decrease in the shReg3a-transduced neurons (Suppl. Figure S1A,B). REG3A depletion led to a modest increase in axon branching (Suppl. Figure S1C). These data raise the possibility that the elevation of axonal *Reg3a* mRNA after PNS injury affects axon growth rates.

**Figure 1:**
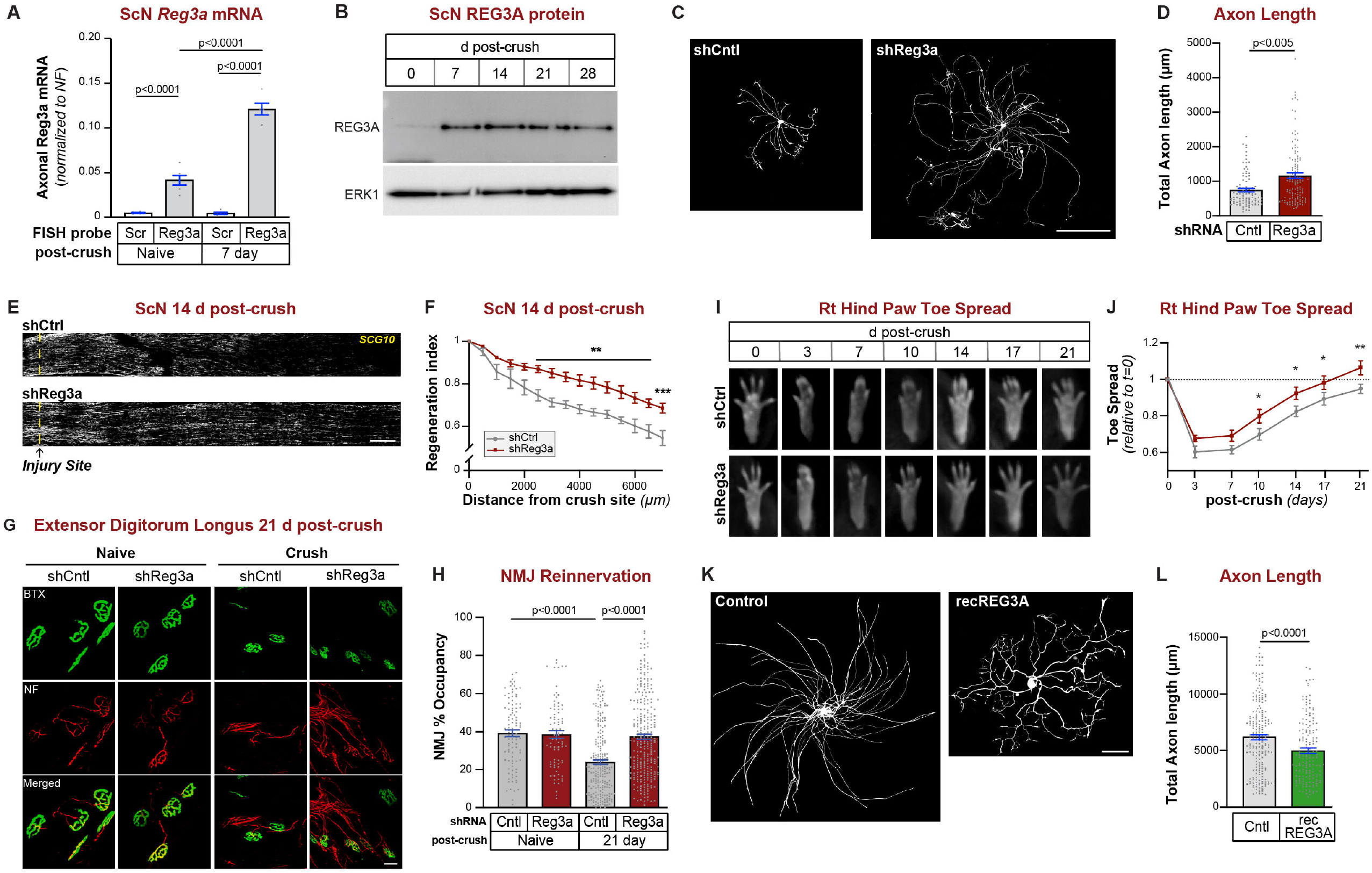
REG3A slows axon growth and regeneration. **A**, Quantitation of smFISH images for naïve vs. 7 d crush injured sciatic nerve (ScN) shows a significant increase in axonal *Reg3a* mRNA. One-way ANOVA with Tukey post-hoc analysis, mean ± SEM (n ≥ 6 mice per group). **B**, Representative immunoblot of 0-28 d post-injury ScN lysates shows increase in REG3A protein signals in regenerating nerve with relatively equivalent levels of ERK1 protein. **C-D**, Representative images for SMI312 (NF) + TuJ1 immunofluorescence in AAV-shCntl vs -shReg3a transduced adult DRG cultures (**C**). Quantitation shows increased axon length with shReg3a vs. shCntl (**D**). See Suppl. Figure S1A-C for shRNA validations and axon branching data. Student’s t-test, mean ± SEM (n ≥ 75 neurons); scale bar = 100 µm. **E-F**, Representative images of exposure matched SCG10 immunostained ScN sections at 14 d post-nerve crush from mice transduced with AAV-shReg3a vs. -shCnt; injury site is marked by dashed line (**E**; left is proximal and right is distal). Regeneration indices for mice as in E (**F**). See Suppl. Figure S1D-J for *in vivo* shRNA validations and regeneration indices on 7, 21 and 28 d post ScN crush. Two-way ANOVA with Sidak post-hoc analysis, mean ± SEM, ^**^ p ≤ 0.01 and ^***^ p ≤ 0.005 (n ≥ 5 per group); scale bar = 500 µm. **G-H**, Representative images of extensor digitorum longus (EDL) neuromuscular junctions (NMJ) from AAV0-shCntl vs. -shReg3a transduced mice at 0 and 21 d post ScN crush injury (**I**). Quantitation of NMJ occupancy is shown in **J**. Two-way ANOVA with Tukey post-hoc analysis (n ≥ 84 NMJs across 6 animals; mean ± SEM; scale bar = 20 µm. **I-J**, Representative images of right (Rt) hind paw (**G**) from BlackBox One analyses of AAV-shRNA transduced mice as in E. AAV-shReg3a transduced mice show increased toe spreading beginning at 10 d post-injury. Quantitation of toe spread is shown in J. See Suppl. Figure S2A-D for data on *KHSRP*^*-/-*^ vs. *KHSRP*^*+/+*^ mice and Suppl. Figure S2E-H for additional behavioral assessments of AAV-shRNA transduced mice wild type mice. Two-way ANOVA with Tukey post-hoc analysis, mean ± SEM, ^*^ p ≤ 0.05 and ^**^ p ≤ 0.005 (n = 10 shCntl and 13 shReg3A). **K-L**, Representative SMI312 + TuJ1 immunofluorescence images for control vs. recREG3A treated adult DRG cultures (**K**). Quantitation of axon morphologies shows decreased length and increased branching after recREG3A exposure (**L**). Student’s t-test, mean ± SEM (n ≥ 163 neurons); scale bar = 100 µm.

### REG3A protein slows nerve regeneration

We used *in vivo* depletion of endogenous *Reg3a* mRNA to determine if REG3A protein slows PNS nerve regeneration. For this, mice were transduced with AAV9-shRNAs by direct injection into the sciatic nerve (shReg3a and shCntl). The shReg3a-transduced mice showed clear *in vivo* reduction in *Reg3a* mRNA and REG3A protein in their axons (Suppl. Figure S1D,E). Sciatic nerve and L4-6 DRGs showed decreased *Reg3a* mRNA by RTddPCR at each time point post-injury with expression of shReg3a (Suppl. Figure S1F,G). The shCntl-transduced mice showed increased *Reg3a* mRNA in the L4-6 DRGs and sciatic nerve following crush injury as we previously reported (8) (Suppl. Figure S1F,G). Nerve regeneration indices increased in the shReg3a-transduced mice, with greatest difference compared to shCntl-transduced mice at 14 d post-injury (Figure 1E,F ; Suppl. Figure S1H-J). The shReg3a-transduced mice also showed increased neuromuscular junction occupancy, suggesting accelerated reinnervation of target tissues following nerve injury (Figure 1G,H).

We next asked whether *Reg3a* depletion affects functional reinnervation of synaptic targets after sciatic nerve injury using the BlackBox Bio system. This instrument measures behaviors of freely moving mice in a dark environment using continuous recording of paw placement and pressure (*9*). To initially test ability to assess nerve regeneration using this approach, we analyzed KHSRP knockout vs. wild type mice, since *KHSRP* deletion accelerates PNS regeneration (*4*). Consistent with previous findings (*10*), *KHSRP*^*-/-*^ mice showed increased motility based on distance traveled before injury compared to wild type mice (Suppl. Figure S2A). The *KHSRP*^*-/-*^ mice further showed accelerated functional recovery after right-sided sciatic nerve crush based on right to left hind paw luminance and right to average front paws luminance compared to *KHSRP*^*+/+*^ mice (Suppl. Figure S2B-D). Since paw luminance showed the most consistent differences at 21 d and earlier after injury in *KHSRP*^*-/-*^ vs. *KHSRP*^*+/+*^ mice, we focused on 0-21 d post-crush interval for assessing recovery within REG3A-depleted mice. Sciatic nerves of wild type mice were bilaterally injected with AAV9-shCntl-BFP vs. -shReg3a-BFP 14 d prior to a right-sided sciatic nerve crush. The AAV-shCntl- and -shReg3a-transduced mice showed no difference in motility (Suppl. Figure S2E); however, right to left hind paw and right hind to average front paws luminance were increased in the shReg3a-vs. shCntl-transduced over 10-21 d post-injury (Suppl. Figure S2F-H). Additionally, right hind paw toe spreading was significantly increased over these same intervals in the shReg3a-vs. shCntl-transduced mice (Figure 1I,J ; note – *KHSRP*^*-/-*^ mice chewed their ipsilateral hind paw toes after crush injury, so toe spreading could not be assessed in this constitutive knockout). Thus, based on both morphological and functional assessments, REG3A depletion accelerates peripheral nerve regeneration.

### REG3A reduces axon regeneration through translational modulation of axonal Khsrp mRNA

The accelerated nerve regeneration seen with REG3A depletion mirrors what we recently published for *KHSRP* knockout mice (*4*). Since *Reg3a* mRNA encodes a lectin-like protein that includes an N-terminal signal peptide for secretion (GenBank ID # NM_011259), we asked if recombinant REG3A (regREG3A) protein affects axon growth. In contrast to effects seen with depleting endogenous REG3A, bath applied recREG3A reduced axon length and increased axon branching in adult DRG cultures (Figure 1K,L; Suppl. Figure 1K). The decrease in axon growth, increase in axon branching, and increase in axonal KHSRP seen with recREG3A application was prevented by the translation inhibitor cycloheximide (CHX; Figure 2A,B; Suppl. Figure S3A-C). We further tested for axonal translation of *Khsrp* using a diffusion-limited GFP reporter mRNA (GFP^MYR^) that includes the 5’ and 3’ UTRs of *Khsrp* mRNA (GFP^MYR^5’/3’*Khsrp*) as a surrogate for the endogenous mRNA (*4*). By fluorescence recovery after photobleaching (FRAP), axonal translation of GFP^MYR^5’/3’*Khsrp* was elevated in response to recREG3A, and this was attenuated by cycloheximide treatment (Figure 2C,D). The axon growth attenuating effects of recREG3A were not seen in DRGs cultured from *KHSRP*^*-/-*^ mice (Suppl. Figure S3D); however, recREG3A treatment still increased axonal calreticulin protein levels in the *KHSRP*^*-/-*^ DRG cultures (Suppl. Figure S3E) and Calreticulin mRNA (*Calr*) is known to be translated in response to elevated axoplasmic Ca^2+^ (*4, 11, 12*). Together, these data suggest that REG3A decreases axon growth by increasing axonal translation of *Khsrp* mRNA, and they raise the possibility that this occurs through Ca^2+^ signaling.

**Figure 2:**
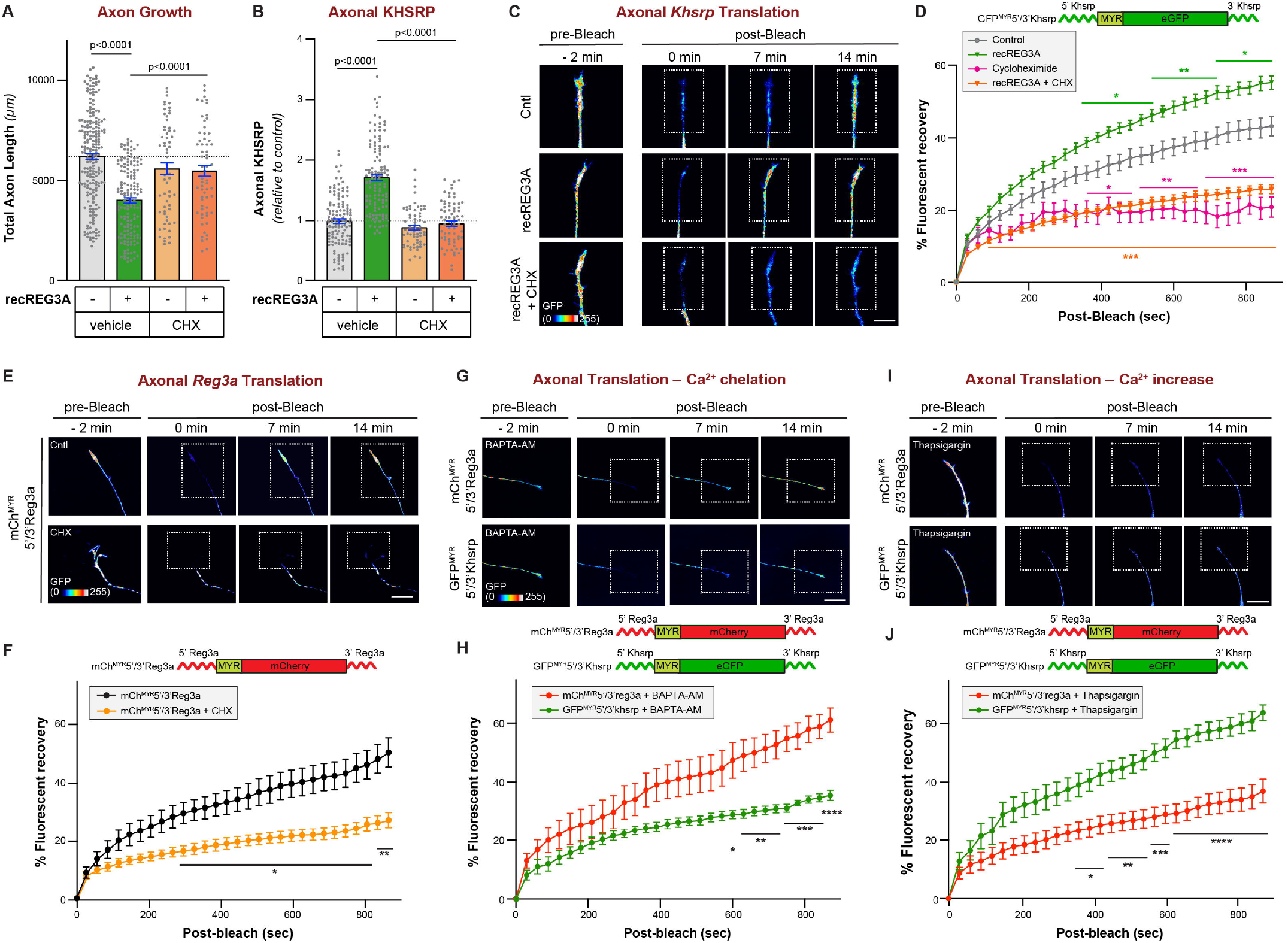
REG3A increases axonal translation of Khsrp mRNA. **A-B**, Axon length quantitation (**A**) and axonal KHSRP (**B**) for DRG neurons treated ± recREG3A and vehicle (DMSO) vs. cycloheximide (CHX). See Suppl. Figure S3A-C for representative images and axon branching data. One-way ANOVA with Tukey post-hoc analysis, mean ± SEM (n ≥ 63); scale bar = 100 µm. **C-D**, Fluorescence recovery after photobleaching (FRAP) for terminal axons of GFP^MYR^5’/3’Khsrp transfected DRGs treated with recREG3A ± CHX shown as representative exposure-matched image sequences (**C**) and quantitation (**D**). Two-way ANOVA with Tukey post-hoc analysis, mean ± SEM, ^*^ p ≤ 0.05, ^**^ p ≤ 0.005, and ^***^ p ≤ 0.0005, with colors matched to dataset vs. control (n ≥ 8 axons across at least three culture preparations); scale bar = 10 µm. **E-F**, Representative exposure-matched image sequences (**E**) and quantitation (**F**) for axonal FRAP in mCh^MYR^5’/3’Reg3a transfected DRGs ± CHX. Repeated measures ANOVA with Tukey post-hoc analysis, ^*^ p ≤ 0.05 and ^**^ p ≤ 0.01, mean ± SEM (n ≥ 10 axons across at least 3 culture preparations); scale bar = 10 µm. **G-J**, Representative exposure-matched image sequences (**G**,**I**) and quantitations (**H**,**J**) for axonal FRAP in mCh^MYR^5’/3’Reg3a + GFP^MYR^5’/3’Khsrp transfected DRGs treated with BAPTA-AM to chelate intracellular Ca^2+^ (**G-H**) or Thapsigargin to increase intracellular Ca^2+^ (**I-J**). Mean ± SEM is shown. Repeated measures ANOVA with Tukey post-hoc analysis, ^*^ p ≤ 0.05, ^**^ p ≤ 0.01, ^***^ p ≤ 0.005, and ^****^ p ≤ 0.00; scale bar = 10 µm.

### Axonal Reg3a and Khsrp mRNAs show opposite translational regulation by Ca^2+^

Consistent with endogenous *Reg3a* mRNA’s localization into axons, a myristoylated mCherry (mCh^MYR^) fluorescent reporter with *Reg3a*’s 5’ and 3’UTRs (mCh^MYR^5’/3’Reg3a) is locally translated in axons of cultured DRG neurons by FRAP analysis (Figure 2 E,F). Since *Khsrp* mRNA translation is increased by elevated axoplasmic Ca^2+^ and decreased by intracellular Ca^2+^ chelation (*4*), we asked if axonal mCh^MYR^5’/3’Reg3a translation is similarly sensitive to axoplasmic Ca^2+^ modulation. In DRG cultures co-expressing mCh^MYR^5’/3’Reg3a and GFP^MYR^5’/3’Khsrp, axonal mCh^MYR^5’/3’Reg3a fluorescent recovery was significantly higher upon exposure to the intracellular Ca^2+^-chelator BAPTA-AM compared to GFP^MYR^5’/3’Khsrp (Figure 2G,H). In contrast, axonal mCh^MYR^5’/3’Reg3a fluorescent recovery was significantly decreased by thapsigargin, which increases cytoplasmic Ca^2+^ levels by blocking Ca^2+^ uptake by the ER, compared to GFP^MYR^5’/3’Khsrp’s (Figure 2I,J). The elevation of GFP^MYR^5’/3’Khsrp translation with thapsigargin and depressed translation of GFP^MYR^5’/3’Khsrp in axons is consistent with previous findings (*4*). Together, these data show that *Reg3a* and *Khsrp* mRNAs are translated under distinctly different intracellular Ca^2+^ concentrations.

### REG3A stimulates release of axonal Ca^2+^ stores and activates an intrinsic stress response

Since *Khsrp* mRNA translation increases with elevated Ca^2+^ and recREG3A increases axonal *Khsrp* translation, we asked if REG3A regulates Ca^2+^ entry or intracellular release. Chelating intracellular but not extracellular Ca^2+^ (using BAPT-AM vs. BAPTA, respectively) blocked the effect of recREG3A on axon outgrowth, axon branching and axonal KHSRP elevation in DRG cultures (Figure 3A,B; Suppl. Figure S3F). By FRAP assay, the recREG3A-dependent increase in axonal translation of *GFP*^*MYR*^*5’/3’Khsrp* was also significantly attenuated by intracellular but not extracellular Ca^2+^ chelation (Figure 3C,D). Thus, REG3A activates release of intracellular Ca^2+^ stores to increase axonal *Khsrp* translation and reduce axon growth.

**Figure 3:**
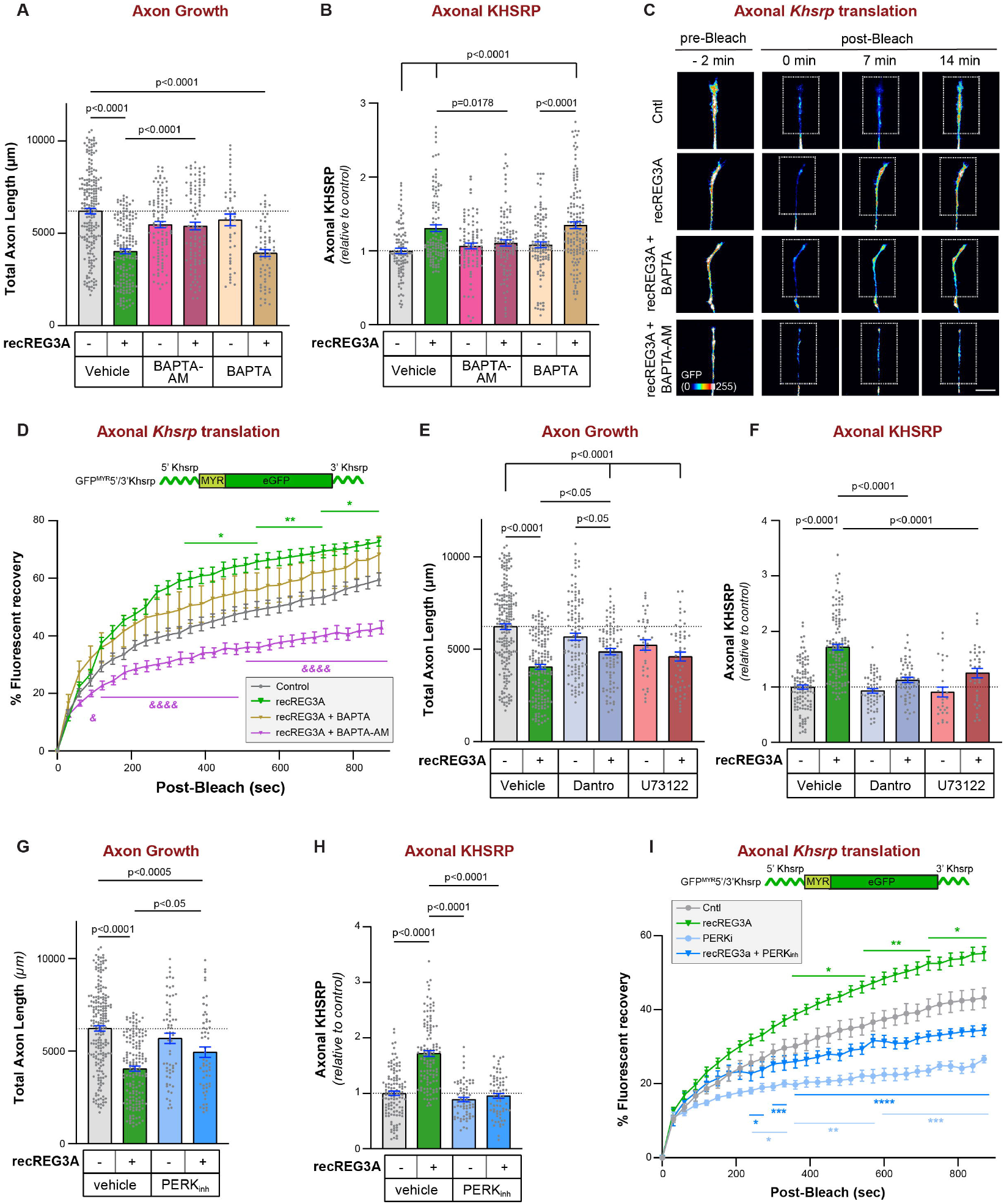
Axonal REG3A signaling requires release of intracellular calcium and an ER stress-like response. **A-B**, Axon growth (**A**) and axonal KHSRP levels (**B**) for DRG neurons ± recREG3A in presence of BAPTA-AM to chelate cytoplasmic Ca^2+^, BAPTA to chelate extracellular Ca^2+^, or vehicle control (DMSO). Suppl. Figure S3F shows axon branching for these cultures. One-way ANOVA with Tukey post-hoc analysis; mean ± SEM (n ≥ 46 in A and ≥ 99 in B). **C-D**, Axonal FRAP analyses are shown for GFP^MYR^5’/3’Khsrp reporter in DRG neurons treated as in A. Representative exposure-matched images sequences shown in **C** and quantitation in **D**. BAPTA-AM attenuates recREG3A-dependent fluorescent recovery of the GFP reporter but BAPTA does not. Two-way ANOVA with Sidak post-hoc analysis; mean ± SEM; ^*^ p ≤ 0.05 and ^**^ p ≤ 0.005 vs. control and & p ≤ 0.05 and &&&& p ≤ 0.001 vs. RecREG3A (n ≥ 10 axons across at least three culture preparations); scale bar = 10 µm. **E-F**, Axon growth (**E**) and axonal KHSRP levels (**F**) for DRG neurons ± recREG3A in presence of dantrolene to inhibit the Ryanodine receptor or U73122 to attenuate production of IP3. Suppl. Figure S3G shows axon branching for these cultures. One-way ANOVA with Tukey post-hoc analysis, mean ± SEM, (n ≥ 36 neurons in E and ≥ 28 in F across ≥ 3 replicate cultures). **G-H**, Axon growth (**G**) and axonal KHSRP levels (**H**) for DRG neurons ± recREG3A in presence of PERK inhibitor (PERK_inh_) vs. vehicle. Suppl. Figure S4A-B shows representative images and axon branching for these cultures. One-way ANOVA with Tukey post-hoc analysis, mean ± SEM (n ≥ 60 in A and ≥ 34 in B). **I**, Axonal FRAP analysis for axonal GFP^MYR^5’/3’Khsrp transfected adult DRG cultures treated as in G. Two-way ANOVA with Tukey post-hoc analysis, mean ± SEM; ^*^ p ≤ 0.05, ^**^ p ≤ 0.005, ^***^ p ≤ 0.0005, ^****^ p < 0.0001 (n ≥ 10 axons across at least three culture preparations with colors matched to condition vs. control).

While ER, mitochondria, and lysosomes are potential sources of Ca^2+^ stored within axons, we had previously found that axonal *Khsrp* translation requires activation of PERK with subsequent phosphorylation of the translation factor eIF2*α* (*4*), a pathway that is activated upon ER Ca^2+^ depletion (*13*). Thus, we asked if inhibition of ER Ca^2+^ channels would impact responses to recREG3A. Inhibition of the ER Ryanodine receptor (RYR) with dantrolene or prevention of ER inositol phosphate-3 receptor (IP3R) activation by inhibition of phospholipase C (PLC) with U73122 each partially reversed recREG3A’s axon growth attenuation and axonal KHSRP elevation (Figure 3E,F; Suppl Figure S3G), suggesting that both RyR and IP3R activation lie downstream of REG3A. Consistent with this, PERK inhibition prevented the reduction in axon growth, increase in axon branching, and increase in axonal KHSRP seen with recREG3A treatment (Figure 3G,H; Suppl. Figure S4A). RecREG3A dependent axonal translation of *GFP*^*MYR*^*5’/3’Khsrp* was prevented by PERK inhibition (Figure 3I). recREG3A stimulation increased axonal eIF2*α*^PS51^ and this was prevented by PERK inhibition (Suppl. Figure S4B-D). Taken together, these data indicate that recREG3A slows axon growth by stimulating Ca^2+^ release from ER stores to activate PERK→eIF2*α*→*Khsrp* translation pathway.

### Endogenous REG3A decreases axon growth by episodically elevating growth cone Ca^2+^

The findings above provided circumstantial but not direct evidence for elevated axoplasmic Ca^2+^ as a key regulator of REG3A’s effects on axons, so we asked whether recREG3A increases Ca^2+^ levels in axons of cultured DRG neurons. DRG cultures transduced with AAV expressing the genetically-encoded Ca^2+^ indicator GCaMP6s showed a clear increase in axonal Ca^2+^ signals after recREG3A stimulation (Figure 4A). Co-transduction with AAV-GCaMP6s plus AAV-shReg3a or -shCntl showed a decrease in basal axonal GCaMP6s signals when endogenous REG3A was depleted (Figure 4B). Axonal signals for the Fluro-4 Ca^2+^ indicator were similarly increased after recREG3A treatment and decreased with AAV-shReg3a-transduction (Suppl. Figure 4E,F). Thus, REG3A can regulate axoplasmic Ca^2+^ levels.

**Figure 4:**
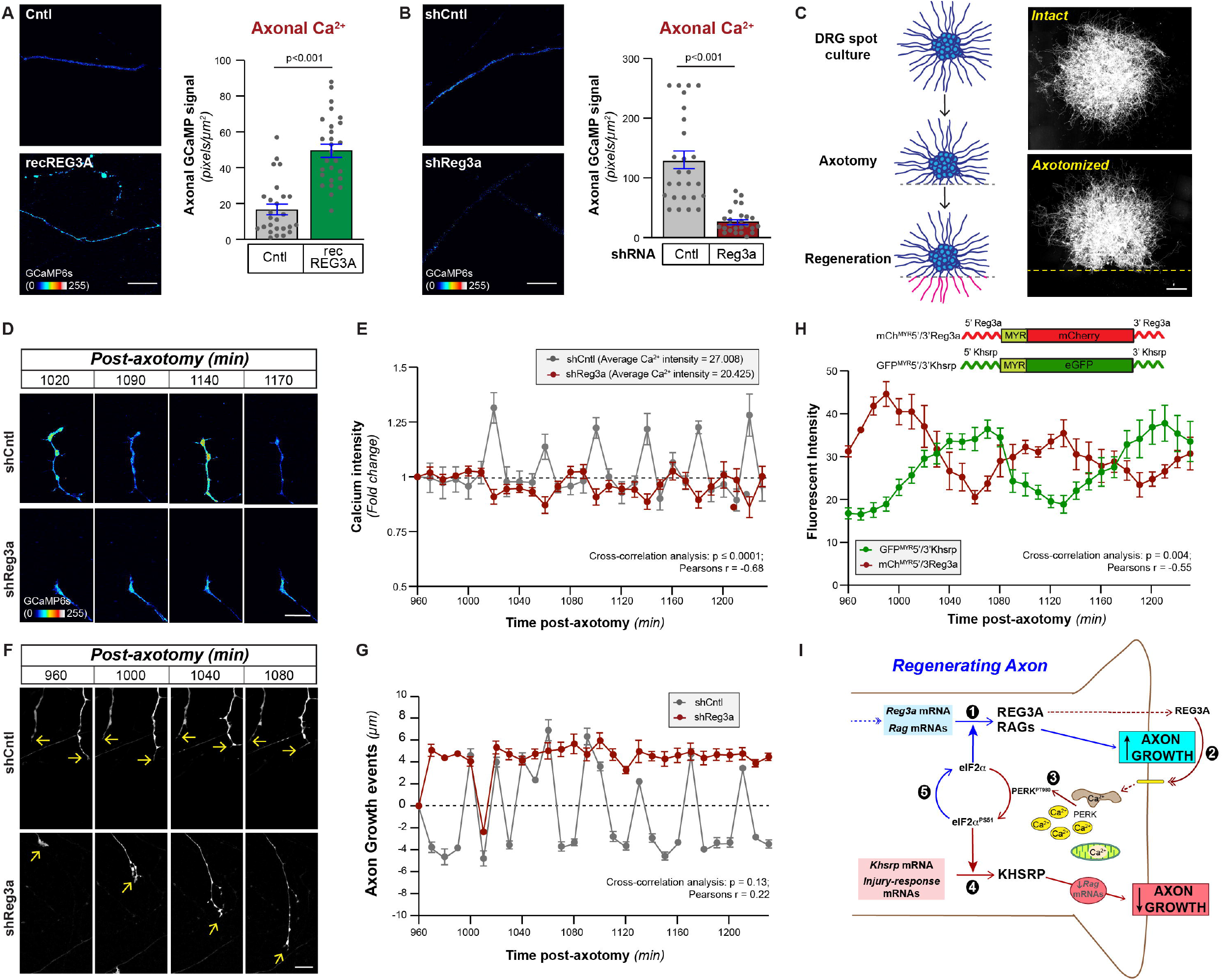
Endogenous REG3A attenuates axon growth by increasing axoplasmic calcium and activating KHSRP synthesis in axons. **A-B**, Representative exposure matched images of intact axons and quantitation for DRGs transduced with AAV-GCaMP6s show elevation of axonal GCaMP6s signals after 1 hr recREG3A treatment (**A**). Representative exposure matched images of intact axons and quantitation for DRGs co-transduced with AAV-shRNAs + AAV-GCaMPS6s show reduction of axonal GCaMP6s signals with shReg3a (**B**). Suppl. Figure S4E-F shows similar responses using the Fura-4 calcium indicator. Imaging parameters for panels A-B and Suppl. Figure S4E-F were set for recREG3A or shReg3A conditions with control parameters matched within each experiment to optimize detecting elevation vs. depression of Ca^2+^ signals. Student’s t-test, mean ± SEM (n ≥ 75 neurons over ≥ 3 culture preparations); scale bar = 10 µm. **C**, Schematic for *in vitro* axotomy in DRG spot cultures with representative images before and after axon severing; scale bar = 100 µm. **D-E**, GCaMP6s signals in distal axons of AAV-shCntl vs. -shReg3a transduced DRG spot cultures are shown in representative exposure matched image sequences (**D**) and quantitation of mean signal intensities ± SEM (**E**) based on average signals across each time point (mean GCaMP6s signals over the entire imaging sequence are approximately 35% higher for shCntl vs. shReg3a-transduced cultures; also see Suppl. Figure S4K). P values by cross-correlation analysis; n = 10 axons across 4 separate culture preparations; scale bar = 10 µm. **F-G**, Axon growth events were assessed in the neurons from panel D based on BFP signals from the AAV-shRNAs. Representative exposure matched image sequences (**F**) and quantitation of axon growth events (**G**) as mean change ± SEM (see Suppl. Figure S4L-M for the overall average events and growth rates across the sequences and Suppl. Video 1). P values by cross-correlation analyses (n = 10 axons across 4 separate culture preparations); scale bar = 10 µm. **H**, Mean growth cone fluorescence ± SEM for DRGs expressing GFP^MYR^5’/3’Khsrp and mCh^MYR^5’/3’Reg3a is shown for proximal severed axons in spot cultures over 16-20.5 h post-axotomy imaging sequences. P value by cross-correlation analyses (n = 10 axons across 4 separate culture preparations). **I**, Schematic for axon-intrinsic translational regulation of *Khsrp* mRNA by axonally synthesized REG3A in regenerating axons. *1*) Injury induces an increase in cell body *Reg3a* mRNA that is transported into regenerating axons over 2-3 d following axotomy; *Reg3a* mRNA is translated under low Ca^2+^ levels when eIF2*α* is not phosphorylated. *2-3*) Nascent REG3A secreted from the distal axon activates a transmembrane signal resulting in release of ER Ca^2+^ to activate PERK resulting in phosphorylation of eIF2*α* on S51. *4*) Increased eIF2*α*^PS51^ activates translation of *Khsrp* mRNA, which promotes decay of regeneration-associated gene (Rag) mRNAs. *5*) Subsequent dephosphorylation of eIF2*α* promotes translation of *Reg3a* to generating an oscillating translation of *Khsrp* and *Reg3a* as shown in panel F.

We next used a DRG ‘spot culture’ technique, where axons are physically and visually isolated from their soma (Figure 4C). Characterizing intact axons in the DRG spot cultures showed increase in both axonal eIF2*α*^PS51^ and KHSRP after recREG3A treatment (Suppl. Figure S4G,H). After manual transection, regenerating proximal axons of segments in these DRG spot culture showed decreased axonal eIF2*α*^PS51^ and KHSRP in AAV-shReg3a-*vs*. shCntl-transduced cultures (Suppl. Figure S4I,J). This axon transection in the spot cultures provided an opportunity to visualize regenerating axons over time. Thus, we used spot culture co-transduction with AAV-GCaMP6s + AAV-shReg3a vs. -shCntl to determine if Ca^2+^ levels dynamically change over time in regenerating axons. Average growth cone GCaMP6s signals were significantly lower in the shReg3a-*vs*. shCntl-transduced neurons beginning at 16 h post axotomy (Suppl. Figure S4K). Fold-changes in Ca^2+^ for each time point relative to the average growth cone GCaMP6s signals across the entire sequence showed peaks in growth cone Ca^2+^ signals with a periodicity of 50-70 min in shCntl-transduced cultures (Figure 4D,E). In contrast, shReg3a-transduced cultures showed significantly less variability in growth cone Ca^2+^ over the imaging sequence (Figure 4D,E). We assessed axon growth events in these same cultures based on the BFP included in the shCntl and shReg3a RNA vectors. Axons of the shCntl-transduced neurons showed alternating extension and retraction events, while the shReg3a-transduced neurons showed overall processive axon growth (Figure 4F,G; Suppl. Figure S4L,M; Suppl. Video S1).

Interestingly, regenerating axons in DRG spot cultures co-transfected with translation reporters showed that fluorescence of GFP^MYR^5’/3’Khsrp and mCh^MYR^5’/3’Reg3a translation reporters exhibited peaks and troughs with opposing periodicity (Figure 4H). Individual axons varied substantially, so this analysis required that the x-axes for individual axons be aligned for an initial peak in the *Reg3a* reporter fluorescence. This variability raised the possibility of an autocrine rather than paracrine response to endogenous REG3A. Consistent with this possibility, adjacent axons in the same field did not show synchronous peaks and troughs for the *Khsrp* and *Reg3a* reporter signals (Suppl. Figure S5A,B). Furthermore, BFP negative axons of non-transduced neurons in the nerves of the AAV-shReg3a-BFP transduced mice in Figure 1E-F showed regeneration indices identical to the BFP postive axons in the AAV-shCntl-GFP-transduced mice (Suppl. Figure S5C).

Immunofluorescent signals for endogenous REG3A protein were increased in cultures of L4-6 DRGs from mice that had previously undergone an *in vivo* sciatic nerve crush for ‘injury-conditioning (*14*) and in regenerating sciatic nerve compared to naïve DRG cultures and uninjured nerve, respectively (Suppl. Figure S5D,E). In both the injury-conditioned DRG cultures and regenerating sciatic nerve, the REG3A signals appeared to be concentrated along the periphery of the axons by orthogonal confocal projections (Suppl. Figure 5SD,E; Suppl. Videos S2 and 3). Axons have functional equivalents of ER and Golgi apparatus for secretion of newly synthesized proteins (*15*). Lectin proteins bind to proteoglycans (*16*), so it is conceivable that secreted REG3A protein adheres to the membrane and juxta-membrane region. Consistent with this, heparinase treatment of DRG cultures prior to immunofluorescent staining drastically decreased the signals for REG3A in the injury-conditioned DRGs (Suppl. Figure S5D, Suppl Video 2). Taken together, these data suggest that axonal synthesized and secreted REG3A signals in an autocrine fashion to slow axon growth through Ca^2+^ store release-dependent translation of *Khsrp* mRNA (Figure 4I).

## Discussion

Many lines of evidence indicate that axonally synthesized proteins provide an initial response to injury and then later promote regeneration of the injured axons through local actions of the newly synthesized proteins (*17, 18*). While most of the axonally-synthesized proteins studied to date promote axon growth or neuronal survival after injury, axonal translation of *Khsrp* impedes axon growth by promoting decay of ARE-containing axonal mRNAs (*4*). *Khsrp* translation is increased by axoplasmic Ca^2+^ elevation following injury, but its protein product stays elevated in axons well beyond the initial axotomy-induced Ca^2+^ elevation subsides (*4*). Here, we show that axonally-synthesized REG3A protein provides a means to intermittently activate *Khsrp* translation in regenerating axons. Consequently, axonal translation of *Reg3a* slows PNS nerve regeneration. The slow rate of PNS nerve regeneration can result in prolonged target tissue denervation and ultimately unsuccessful recovery from injury for lesions requiring axon growth of more than a few centimeters to reach target tissues (*3*). Thus, interventions that can accelerate axon regeneration have great potential to increase recovery from PNS nerve injuries, and our data point to an axon-intrinsic REG3A→Ca^2+^→KHSRP pathway as a target for accelerating nerve regeneration (Figure 4I).

REG3A is a member of a family of lectin-like proteins that includes REG3B and REG3G. Murine REG3A is about 59% identical to REG3B and REG3G proteins. REG3 proteins have been functionally linked to regeneration of pancreatic islet cells and REG3A has bactericidal activity in the intestine (*7*). EXTL3 protein was reported as a cell surface receptor for REG3 proteins (*19*). Several inflammatory cytokines have been reported to increase REG3A expression in epithelial systems where REG3A binding to EXTL3 activates the phosphatidyl-inositol-3 kinase (PI3K) to AKT signaling pathway (*20*). REG3G has also been reported to activate PI3K-AKT as well as the RAS/RAF-ERK signaling pathways through EXTL3 binding (*21-23*). Though EXTL3 has well-established roles as a glycosyl-transferase in the Golgi apparatus (*24*), cell surface localization of EXTL3 has been reported for hippocampal neurons, N2a cells and neuron-like PC12 cells (*25*). Further, REG1*α* protein, which shares less homology to REG3A than the REG3 family members (42% sequence identity), was reported to promote neurite growth in PC12 cells by binding to EXTL3 (*25*). PI3K, AKT, Ras/Raf and ERK signaling are typically growth-promoting in neuronal contexts rather than the clear attenuation of growth that we see with both recombinant and endogenous REG3A in the experiments here. REG3B was also shown to activate JAK/STAT3 signaling in carcinoma cells (*26*), but STAT3 activation after injury promotes nerve regeneration and neuronal survival (*27*). Thus, REG3A’s attenuation of axon growth seems unlikely to occur through previously defined signaling events downstream of EXTL3 binding.

REG3A’s function as a bactericide is linked to its glycoprotein-binding activity, with hexamers of REG3A protein dimers permeabilizing the bacterial membrane (*28, 29*). Cell membrane permeabilization by a naturally-occurring peptide fragment of REG3A (termed PAP) has also been reported to stimulate pancreatic beta-cell proliferation (*30*). Pore-forming activities reported for REG3A could promote Ca^2+^ entry after REG3A secretion or recREG3A exposure, but we find that neither the axon growth attenuation nor the axonal KHSRP elevation by REG3A was affected by depletion of extracellular Ca^2+^. Activation of PERK and subsequent eIF2*α* phosphorylation increasing axonal KHSRP synthesis is consistent with release of ER Ca^2+^ stores in response to both recombinant and endogenous REG3A. Though Ca^2+^ influx can trigger subsequent ER Ca^2+^ release (*31*), extracellular Ca^2+^ chelation did not prevent effects of recREG3A on DRG axons, so REG3A-dependent Ca^2+^ entry cannot explain the effects of REG3A seen here. Taken together, our data argue that the effects of axonally-synthesized REG3A are through a different receptor and signaling mechanism than previously published for REG3A. Further our data suggest that this is an autocrine, axon-intrinsic mechanism that slows axon growth.

Several lines of evidence indicate that axotomy triggers an unfolded protein or integrated stress response locally that increases eIF2*α*^PS51^ levels to impact translation in axons (*6, 11, 32-34*). Consistent with this, we see that recREG3A increases axonal levels of calreticulin protein, whose mRNA is translated in axons by elevated Ca^2+^ and PERK activation (*6, 11, 12*). Inflammation and injury have been reported to increase REG3A expression in rodent sensory neurons within 2 d following the inflammation or injury (*35*). Although the authors did not address function of the neuronal REG3A, they concluded that increased peripheral nerve REG3A levels derive from anterograde transport based on nerve ligation studies (*35*). Axonal localization of *Reg3a*, combined with localized translation of an axonal reporter mRNAs with *Reg3a*’s 5’ and 3’UTRs indicate that the increases in axonal REG3A protein after axotomy derive from anterograde mRNA transport and subsequent translation in axons. Together, these observations suggest that targeting the signaling mechanisms underlying this REG3A to KHSRP signaling can bring new strategies for accelerating nerve regeneration.

## ACKNOWLEDGEMENTS

JLT is the incipient University of South Carolina SmartState Chair in Childhood Neurotherapeutics.

## AUTHOR CONTRIBUTIONS

CNB, JYL, LSV, SM, AK, and JLT designed experiments.

CNB, JYL, LSV, SM, MCH, MC, LFT, AM, IDC, and MLDL performed experiments.

CNB, JYL, LSV, SM, MCH, MC, LFT, and AM provided data analyses.

CNB, JYL, SM, MCH, MC, LFT, AM, MLDL, and ET performed animal husbandry and mouse colony maintenance.

LSV and JLT supervised the work.

CNB, JYL, LSV, AK, NPB, and JLT wrote and edited the manuscript.

NPB and JLT obtained grants funding the work.

## DATA AND MATERIALS AVAILABILITY

Data included here will be submitted to Zenudo online publically accessible database once accepted for publication. Materials will be freely shared subject to Univ SC policy for Materials Transfer Agreement.

## Supplementary Materials

### MATERIALS AND METHODS

#### Key Reagents and Resources

Supplementary Table S1 summarizes key reagents utilized in these studies and the sources of those reagents where indicated.

#### Animal Use

The Institutional Animal Care and Use Committee of the University of South Carolina approved all animal procedures. Adult (8-16 wks old) male and female *Khsrp* knockout (*Khsrp*^*-/-*^) (*36*) and wildtype (C57Bl/6) mice were used for all experiments. Isoflurane inhalation was used for anesthesia in all survival surgery experiments (see below) and animals were euthanized by CO_2_ asphyxiation as indicated for results.

For peripheral nerve injury, 8-12 week old, anesthetized mice were subjected to sciatic nerve crush at mid-thigh level as previous described (*4*). Briefly, the sciatic nerve was exposed crushed using #2 fine jeweler’s forceps twice for 15 sec each, ∼1.5-2 cm from its origin, proximal to the trifurcation. Axotomy was monitored by the initial contraction of the hind limb upon application of pressure to the sciatic nerve and then the lack of hind paw extension during and upon recovery from anesthesia. For animals undergoing unilateral nerve crush, the contralateral nerve was exposed but not crushed (*i.e*., ‘sham’ control).

We have previously shown that nerve injected AAV is retrogradely transported to the neuronal cell bodies whose axons transverse the nerve resulting in a neuronal specific transduction (*4*). For shRNA mediated knockdown experiments 4 µl of 5 × 10^11^ particles of AAV9-TagBFP2-U6-shReg3a or TagBFP2-U6-scramble-shRNA (Vector Builder, Chicago, IL) were diluted in 600 mM NaCl (total volume = 5 µl) and injected into the proximal sciatic nerve at ∼1.2-1.5 cm from origin. 14 d after viral transduction, animals were subjected to a bilateral sciatic nerve crush at ∼0.5 cm distal to injection site on both the left and right sciatic nerve as above. For consistency between animals, a single experimenter performed the viral infections and crush injuries within each series of animals.

#### Cell Culture

Adult mouse dorsal root ganglia (DRG) were harvested and placed in ice-cold Hibernate-A medium (BrainBits). For experiments with naïve DRG neurons, all DRGs including lumbar, thoracic and lower cervical were collected. For *in vivo* injury conditioning, only lumber segment 4-6 DRGs were collected. DRGs were rinsed five times with DMEM/F12 + 10U/mL Pen/Strep (CAT#15-140-122, Fisher, Waltham, MA) and then ganglia were treated with 0.125U/mL collagenase type II CAT#17-101-015, Fisher) in DMEM/F12 for 30 min at 37°C, 5 % CO_2_. DRGs were triturated using a fire polished Pasteur pipette and pelleted by centrifuged at 100 xg for 10 min. DRGs were washed twice by resuspension with DMEM/F-12 + Pen/Strep and centrifugation.

Following washes, cells were resuspended in DMEM/F-12 media supplemented with 10 % fetal bovine serum (Gibco, #10437028) 10 µM cytosine arabinoside (Sigma, St Louis, MO; # C6645), 1x L-glutamine (Fisher; # A2916801), 1x N1 medium supplement (0.5 mg/mL Insulin (Sigma; # I9278), 0.5 mg/mL Sodium Selenite (Sigma; # S5261), 0.5 mg/mL Transferrin (Sigma; #T8158), 1.6 mg/mL Putrescine (Spectrum; # P1834), 0.73 ug/mL Progesterone (Spectrum; # P1834), in Earle’s Buffered Salt solution, no phenol (Thermo Scientific, AAJ67559AE). Dissociated DRGs were plated immediately or transfected and then plated (see below) on poly-L-lysine/laminin-coated glass substrates. For coating substrates, coverslips were incubated in 50 µg/ml poly-L-lysine (Fisher; #A005C) for 2 h at 37°C and 5 µg/ml laminin (Sigma) at 4°C overnight. 12 mm glass coverslips, 35 mm black wall glass bottom dishes (WillCo Wells from Ted Paella, Redding, CA; # HBSB-3522), or chambered coverslips (Ibidi, Gräfelfing, Germany; # 80297) were used.

For DRG ‘spot cultures’, isolated and ganglia were treated with collagenase as above for 30 min at 37°C, 5% CO_2_. DRGs were then triturated 10-15 times with a 1000 µl pipette tip followed by a second incubation at 37°C, 5% CO_2_ for 10 min. Ganglia were dissociated into single cell suspension using a 1000 µl pipette tip in 4 ml of media containing 1x penicillin/streptomycin solution. Dissociated ganglia were centrifuged for 10 min at 700 xg. Cell pellet was resuspended in fresh DRG culture medium and plated at a density of approximately 2.8 DRGs per 7 µl (up to 45 DRGs harvested per mouse). For this, dissociated ganglia were gently triturated 10 times with 20 µl pipette tip and 7 µl placed onto poly-D-lysine/laminin-coated chambered coverslips at up to 2 well-separated ‘spots’ per well. Spotted DRGs were incubated at 37°C. 5% CO_2_ for 7 min and then DRG culture medium was added along the wall of the culture vessel adequate to fully cover the cells and not be at risk for evaporation. These ‘spot cultures’ were incubated at 37°C, 5% CO_2_ for 7-8 d. For medium changes, half volume of the medium in each well was removed and replenished with fresh DRG culture medium at d 1 and 5 post-dissociation. To axotomize the DRG spot cultures for *in vitro* regeneration assay, axons were cut on one side of the spot under a stereomicroscope using a sterilized 0.6 mm thick × 2.75 mm wide flat carbon steel surgical blade (Fine Science Tools, 10035-10). Axons were cut at distance approximately equal to the radius of the spot (*i.e*., approximately 1 mm from the closest soma). Axon regrowth from the injury site was evaluated at 16-30 h post-axotomy as indicated in the results section.

For transducing spot cultures with AAV9, dissociated ganglia resuspended in DRG media were pelleted by centrifugation at 700 xg for 10 min. The pellet was resuspended in fresh DRG culture media containing 5 µl of 5 × 10^11^ particles AAV9shReg3a-TagBFP2 or AAV9-shControl-TagBFP2 virus.

For cDNA transfections, dissociated ganglia were pelleted after washing in DMEM/F-12 at 100 x g for 5 min and then resuspended in 100 µl ‘nucleofector solution’ for Rat Neuron Nucleofector kit (Lonza, Alpharetta, GA; # VPG-1003) or 20 µl solution for *Small Cell Number-SCN kit* (Lonza; # VSPI-1003). 4-6 µg plasmid was electroporated for the Rat Neuron Kit and 1 µg for the Small Cell Number Kit using AMAXA Nucleofector apparatus (program G-013 for Rat Neuron kit and SCN-8 for SCN kit). Transfected DRGs were then plated as above.

For stimulation and inhibition in DRG cultures the following agents were used: recombinant human REG3A-His (277.8 nM; SinoBiological, Wayne, PA; # 11235-H08H), cycloheximide (150 µg/ml, Sigma), BAPTA-AM (3 µM; Sigma, St Louis, MO), BAPTA (3 µM; Cayman Chemical, Ann Arbor, MI), Thapsigargin (1 µM; Sigma), GSK260614 (90 µM; Bio-Techne Corp/Tocris, Minneapolis, MN), Dantrolene (10µM, Sigma, # D9175), and U73122 (1 µM; Sigma, # 1268) 36 h after initial plating. Equal concentration of vehicle was used for inhibitors and chelators, and equivalent concentration of BSA was used for recREG3A. For chelators and inhibitors, cultures were incubated for 15 min prior to other manipulations. Cultures were then treated with recREG3A or control for 24 h followed by rinsing in 1 x PBS and fixation in 4% paraformaldehyde in 1x PBS.

#### Plasmid Constructs and virus constructs

All Fluorescent reporter constructs for analysis of RNA translation were based on eGFP with myristoylation element (GFP^myr^; originally provided by Dr. Erin Schuman, Max Planck Institute) or mCherry plasmid with myristoylation element (mCh^myr^). Reporter construct containing 5’ and 3’UTR of *Khsrp* mRNA has been published (GFP^MYR^5’/3’Khsrp) (*4*). For Reg3a translation reporter (mCh^MYR^5’/3’Reg3a), the 5’ and 3’UTRs of mouse *Reg3a* were PCR-amplified and subcloned into mCh^myr^ reporter. pAAV-hSynapsin1-GCaMP6s-P2A-mRuby3 Ca^2+^ sensor plasmid was originally generated by Lin Tian and was purchased from Addgene (*37*). AAV5 and AAV9 preparations were purchased directly from Addgene (112005-AAV5 and 112005-AAV9). AAV9-shCntl-BFP and -Reg3a-BFP were purchased from VectorBuilder.

#### RNA isolation and analyses

RNA was isolated from cultured DRG neurons and sciatic nerves using the *RNeasy Microisolation kit* (Qiagen, Hilden, Germany). Dissociated DRG cultures were grown for 3 d, washed briefly with PBS then RNA was isolated following manufacturer’s protocol. Sciatic nerves were cut into small pieces and digested with collagenase at 37°C for 30 min with intermittent trituration. RNA was isolated from the collagenase-treated nerves using the *RNAeasy Microisolation kit*. RNA yield was quantified using fluorimetry with *Ribogreen* reagent (ThermoFisher, Waltham, MA) and 50 ng of RNA was reverse transcribed using the *Sensifast cDNA synthesis kit* (Bioline, London, UK). Droplet digital (dd) PCR products were detected using Evagreen reagent on a QX200 droplet reader (Biorad, Hercules, CA). Mitochondrial 12S RNA (*Mtrnr1*) mRNA levels were used for normalizing yields across different isolates as indicated in results. The following primers were used for ddPCR (all from Integrated DNA Technologies [IDT], Coralville, Iowa; all listed as 5’ to 3’): *Mtrnr1*, sense – GGCTACACCTTGACCTAACG and antisense – CCTTACCCCTTCTCGCTAATTC; *Reg3a*, sense – TCTACAAGAGAGACAAGATGCTG and antisense – AGCTGGTACGTGGAGAGG.

#### Single Molecule fluorescence in-situ hybridization (smFISH)

smFISH plus immunofluorescence (IF) was used to detect *Khsrp* and *Reg3a* mRNAs in dissociated DRG cultures and sciatic nerve sections. Custom designed Cy3- and Cy5-labelled Stellaris probes (LGC Biosearch Tech, Middlesex, UK) for mouse *Khsrp* and *Reg3a* with Cy3- and Cy5-labelled scramble probes for control were used. Primary antibodies for smFISH/IF consisted of axon marker antibody containing mouse anti-b3-Tubulin (TUJ1, Biolegend; # 801201) and anti-Neurofilament marker (SMI312; Biolegend; # 837904) cocktail (1:200 for tissues and 1:500 for dissociated cultures) and chicken anti-GFP (1:200 for tissues and 1:500 for dissociated cultures; Abcam, Boston, MA; # ab13970). FITC-conjugated donkey anti-mouse and Alexa405-conjugated anti-chicken (1:500 each; Jackson ImmunoRes., West Grove, PA) were used as secondary antibodies.

For cultured neurons, smFISH/IF was performed as previously described with minor modifications (8). All steps were carried out at room temperature and all solutions were RNase free and made using DEPC-treated water unless otherwise noted. Coverslips were briefly rinsed in 1 x PBS and then fixed in 2% PFA in 1 x phosphate-buffered saline (1x PBS) for 15 min. Coverslips were rinsed 2 times in 1x PBS, then permeabilized in 0.3% Triton X-100 in 1x PBS for 10 min. Samples were prehybridized for 1 h in hybridization buffer (50% dextran sulphate, 10 µg/ml *E. coli* tRNA, 10 mM ribonucleoside vanadyl complex, 80 µg BSA, and 10% formamide in 2x SSC), and then incubated with 12.5 µM each Stellaris probe, mouse anti-b3-Tubulin (TUJ1) and anti-Neurofilament marker (SMI312) cocktail (1:500 each), and chicken anti-GFP (1:500) in hybridization buffer for 16 h at 37°C. Coverslips were then washed in PBS + 0.3% Triton X-100 3 times, followed by incubation with FITC-conjugated donkey anti-mouse and Alexa405-conjugated donkey anti-chicken for 1 h. After rinse in PBS, cells were post fixed in 2% PFA in 1x PBS for 15 min and washed 3 times with 1x PBS then DEPC-treated water. Coverslips were inverted and mounted on glass slides using Prolong Gold Antifade (ThermoFisher).

For sciatic nerve tissue, smFISH/IF was performed as previously described with minor modifications (8). All steps were carried out at room temperature and all solutions were RNase free and made using DEPC-treated water unless otherwise noted. Sciatic nerve segments were fixed for overnight at 4°C in 4% PFA, washed with PBS then cryoprotected overnight in 30% buffered sucrose at 4°C. Nerves were cryosectioned at 20 µm thickness and stored at -20°C until used). Sections were brought to room temperature, washed three times in 1 x PBS for 5 min each, and then incubated with 20 mM glycine followed by 0.25 M NaBH_4_ in PBS (3 times, 10 min each for both) to quench autofluorescence. Sections were quickly rinsed in 0.1 M Triethylamine (TEA) and then incubated in 0.1 M TEA + 0.25% acetic anhydride for 10 min. After washing in 2x SSC, sections were permeabilized with 0.3% Triton X-100 in PBS for 10 min. Sections were rinsed with 1x PBS and incubated with 2x SSC + 10% formamide for 10 min. Sections were prehybridized for 1 h in hybridization buffer (50% dextran sulphate, 10 µg/ml E. coli tRNA, salmon sperm DNA, 10 mM ribonucleoside vanadyl complex, 80 µg BSA, and 10% formamide in 2x SSC), and then incubated with 12.5 µM each Stellaris probe in addition to mouse anti-b3-Tubulin (TUJ1) and anti-Neurofilament marker (SMI312) cocktail (1:200 each), and chicken anti-GFP (1:200) in hybridization buffer for 16 h at 37°C. The following day, sections were washed in 2x SSC + 10% formamide at 37°C twice for 30 min each, followed by a 10 min incubation in 0.5x SSC at 37°C. Sections were briefly rinsed in 1x PBS + 1% Triton-X100 and incubated with FITC-conjugated donkey anti-mouse and Alexa405-conjugated donkey anti-chicken for 1 h. Sections were washed in three times in 1x PBS and DEPC-treated water, then mounted under glass coverslips with Prolong Gold.

smFISH and IF signals in tissue sections were imaged using Leica SP8X or Stellaris confocal microscope. 63x/NA 1.4 oil immersion objective and pulsed white light laser was used for imaging RNA in both culture and tissue samples. Scramble probe was used to set the maximum image acquisition parameters (laser power, HyD or PMT detector energy, & offset) that gave minimum signals with the control probes. XYZ image stacks were acquired across at least three separate locations in each section scanned nerve sections.

#### Immunofluorescence (IF)

Standard IF was performed as previously described with all steps at room temperatures unless specified otherwise. Coverslips were fixed with 4% paraformaldehyde in 1 x PBS for 15 min at room temperature and washed 3 times in 1 x PBS. PBS washed neurons were permeabilized with 0.3% Triton X-100 in PBS for 15 min and then placed in block buffer (1X PBS + 10% Normal Donkey Serum [Jackson ImmunoRes]) for 1 h. Neurons were then incubated with primary antibodies diluted in block buffer overnight in a humidified chamber at 4°C. Primary antibodies consisted of chicken anti-NFH/-NFM/-NFL cocktail (1: 500; Aves Lab, Tigard, OR; NFH # AB2313552, NFM # AB2313554, and NFL # AB2313553), mouse anti-b3-Tubulin (TUJ1, Bio Legends, 1:500), anti-Neurofilament marker (SMI312, Bio Legends, 1:500), rabbit anti-KHSRP (1:500; Novus Biologicals, Centennial, CO; #NBP1-18910), rabbit anti-REG3A (1:200; ThermoFisher; PA5-76091), chicken anti-GFP (1:500; Abcam), mouse anti-eIF2*α* (1:200; Cell Signaling Tech., Danvers, MA; 2103S), and rabbit anti-eIF2*α*^PS51^ (1:200; Cell Signaling Tech.; 9721S). After washes in 1x PBS, coverslips were incubated with a secondary antibody cocktail containing FITC-conjugated donkey anti-chicken, Cy3-conjugated donkey anti-mouse, or Cy5 conjugated donkey anti-rabbit diluted in block buffer (all at 1:500; Jackson ImmunoRes., West Grove, PA) for 1 h at room temperature. Coverslips were then washed 3 times in 1x PBS, rinsed with distilled H_2_O, and mounted with Prolong Gold.

For regeneration studies on mouse sciatic nerve and quantifying axonal content of KHSRP and REG3A *in vivo*, sciatic nerve segments were fixed, cryoprotected, cryosectioned, and stored as above. Tissue sections were brought to room temperature, equilibrated in 1x PBS and incubated with 20 mM Glycine for 30 min followed by 0.25 NaBH_4_ for 30 min to quench autofluorescence. Sections were permeabilized in 0.3% Triton X-100 in 1x PBS for 15 min and blocked for 1 h at room temperature in 1x PBS containing 20 mM Glycine, 0.1% Triton X-100, and 10% Normal Donkey Serum. Primary antibodies consisted of, rabbit anti-KHSRP (1:200; Novus Biologicals, Centennial, CO; #NBP1-18910), rabbit anti-REG3A (1:200; ThermoFisher PA5-76091), rabbit anti-Stathmin-2/SCG10 (1:200; Novus Biologicals; #NBP1-49461), mouse anti-b3-Tubulin (TUJ1, Bio Legends, 1:200), anti-Neurofilament marker (SMI312, Bio Legends 1:200) (1:500 each), and chicken anti-GFP (Abcam, 1:200). The GFP antibody cross-reacts with BFP, so it was used to detect BFP signals from AAV-shRNA transduction. Samples were incubated overnight with primary antibodies in a humidified chamber at 4°C. The following day, samples were washed twice with 1X PBS and incubated with secondary antibodies consisted of FITC-conjugated donkey anti-chicken, Cy3-conjugated donkey anti-mouse, and Cy5-conjugated donkey anti-rabbit diluted in block buffer (all at 1:500; Jackson ImmunoRes.) for 1 h at room temperature. Sections were washed with three times with 1X PBS and once with H_2_O prior to mounting with Prolong Gold Antifade.

For DRG spot cultures, chamber wells were gently rinsed with PBS (37°C) and then fixed in 4% paraformaldehyde in PBS for 30 min at room temperature. After additional rinse in 1x PBS at room temperature, wells were incubated with primary and secondary antibodies as outlined above. Following this, chambers were removed and coverslips were mounted on glass slides using Prolong Gold.

All samples were mounted using *Prolong Gold Antifade* and samples were visualized using epifluorescent or confocal microscopy. Leica DMI6000 epifluorescent microscope with ORCA Flash ER CCD camera (Hamamatsu) was used for epifluorescent imaging. Confocal imaging for immunofluorescence was performed on a Leica SP8X microscope fitted with a galvanometer Z stage and HyD detectors; HC PL Apo 63x/1.4 NA objective (oil immersion) was used with acquisition parameters matched for individual experiments using LASX software. Z-stack images were post-processed by Leica *Lightning Deconvolutio*n integrated into LASX software. Deconvolved image stacks were projected into single plane images.

#### Fluorescent recovery after photobleaching (FRAP)

FRAP was performed using a Leica confocal microscope fitted with an environmental chamber to maintain living cells at 37°C and 5% CO_2_. A 63x/NA 1.4 oil immersion objective was used for imaging with the pinhole set to 3 Airy units (AU) for pre-bleach, bleach, and post-bleach sequences to ensure entire thickness of axons was exposed to laser emission (*38*). Dissociated mouse DRG cultures transfected with GFP^MYR^5’/3’khsrp and/or mCherry^MYR^5’/3’reg3a, as indicated, were equilibrated in complete culture media that excluded phenol red. 72 h following transfection, GFP and/or mCherry expressing neurons were chosen for FRAP. A region of interest (ROI) in the most distal portion of the axon (40 x 40 µm, ≥ 250 µm from the soma) was photobleached with 488 nm (GFP) and/or 555 nm (mCherry) argon laser set at 100% power for 80 frames at 0.78 sec each. Pre-bleach and post-bleach signals were captured using 70% power for 488 and/or 555 nm laser line every 30 sec (2 images taken pre-bleach, and 30 images for post-bleach). Translation dependence of recovery was tested by pre-treating DRG cultures with 150 µg/ml cycloheximide (Sigma) 15 min prior to photobleaching. For testing recREG3A- and Ca^2+^-dependent translation by FRAP, transfected DRG cultures were pretreated for 15 min with 5 µg/ml recREG3A, 3 µM BAPTA-AM, 3 µM BAPTA, or 90 µM GSK260614. For combined treatments with recREG3A, DRG cultures were pretreated with inhibitors/chelators 15 min prior to recREG3A treatment.

#### Immunoblotting

DRG cultures were lysed, or sciatic nerves were minced and lysed in RIPA buffer (50 mM Tris-HCl [pH 8.0], 1% NP-40, 0.5% Sodium Deoxycholate, 0.1% SDS, 150 mM NaCl) plus protease inhibitors (Thermo Scientific A32965). Lysates were centrifuged at 20,000 x g for 15⍰min at 4°C. Protein concentrations of supernatants were determined using Pierce BCA Protein Assay Kit (ThermoFisher; # 23227). After normalization for protein content, lysates were denatured in Laemmli sample buffer (62.5 mM Tris [pH 6.8], 2% SDS, 10% glycerol, 1% β-mercaptoethanol, 0.01% bromophenol blue) at 95°C for 5⍰min followed by fractionation on standard SDS/PAGE. Fractionated proteins were electrophoretically transferred to PVDF membranes (GE Healthcare Life Sciences, Marlborough, MA). Membranes were blocked in 5% non-fat dried milk powder (BioRad, Hercules, CA) in Tris-buffered saline (TBS) with 0.1% Tween-20 (TBST) for 1 h at room temperature. Blots were probes overnight at 4°C with the following antibodies diluted in TBST plus 3% BSA: rabbit anti-REG3A (Invitrogen; # PA5-76091), rabbit anti-ERK1 (Abcam; # ab109282), or mouse anti-HIS (Abcam; # ab18184). Membranes were washed in TBST and then incubated with horseradish peroxidase (HRP)-conjugated anti-rabbit IgG (1:2000; Cell Signaling Tech.) diluted in blocking buffer for 1 h at room temperature. Blots were washed in TBST and signals were detected by ECL Prime™ (GE Healthcare Life Sciences, Marlborough, MA).

#### Behavioral analyses of axonal regeneration

Behavioral testing was conducted using adult male and female C57BL/6J mice, including both *KHSRP*^*-/-*^ and *KHSRP*^*+/+*^ genotypes that were age- and sex-matched. Mice underwent a unilateral sciatic nerve crush injury, with corresponding sham surgery to the contralateral side on d 0, and behavioral recordings were acquired at d 0 prior to sciatic nerve crush and every 3-4 d up to 28 d post-crush. A separate cohort of C57BL/6J mice received a bilateral sciatic nerve injection of either shCtrl or shReg3a on d 0. On d 14 post-injection, these animals were subjected to a unilateral sciatic nerve crush, with a contralateral sham surgery. Behavioral recordings for this cohort were taken at d 0, prior to injection, and then every 3-4 d up to 21 d post-crush (35 d post-injection).

All mice were housed in standard clear plastic cages under controlled conditions (temperature 20-26°C, humidity 30-70%, lights on 07:00-19:00) throughout the duration of experiments and all recordings were done between 10:00 and 17:00 in the same room where the animals were housed using the BlackBox One instrument (BlackBox Bio, Cambridge, MA US). The animal containment chamber consisted of an 18 (l) × 18 (w) × 15 cm (h) black acrylic box that was closed on 4 sides and top; bottom of the box consisted of 5 mm thick borosilicate float glass for video recordings with 850 nm near-infrared (NIR) LED strips aligned perpendicular to 2 opposing edges of the glass and 2 separate 850-nm NIR LED strips positioned horizontally 10 cm below the glass floor. These NIR LEDs provide illumination of the animals for camera mounted beneath to recording paw placement/usage and animal movement as described (*9*).

Animals were placed into individual chambers within the device and allowed 10 min of habituation prior to each recording. After 5-10 min habituation period, animals were briefly removed to clean the glass surface of urine, feces, and any debris and then returned to individual chambers for a 20 min recording.

#### Image analyses and processing

*ImageJ* was used to quantify protein and RNA levels in sciatic nerve tissues from optical planes of XYZ scans (*39*). Axon only signals were extracted via *Colocalization plug-in* that to project protein or RNA signals that overlap with axonal markers in each plane to a separate channel. These ‘axon only’ signals were then quantified in each XY plane of these axon only channels. Axon marker signal area was used to normalize signal intensities across the individual XY planes. The relative signal intensity was then averaged for all tiles in each biological replicate.

To assess regeneration *in vivo*, tile scans were post-processed by *Straighten plugin for ImageJ* (http://imagej.nih.gov/ij/). SCG10 fluorescence intensity was measured along the length of the nerve using ImageJ. Regeneration index was calculated by measuring the average SCG10 intensity in 500 µm bins across the length of the section starting at the crush site. The crush site was defined by the position along the nerve length with maximal SCG10 intensity (secondarily confirmed by DAPI signals and DIC images). Dissociated DRGs were immunostained with neurofilament antibodies as described above and axonal morphology was analyzed from epifluorescent tile scans using *WIS-Neuromath* to provide length and branch density information for each neuron (*40*).

For FRAP image sequences, raw images were analyzed for recovery in the bleached ROI using Leica Confocal Software. Percent fluorescent recovery was determined relative to pre-bleach and post-bleach signals, which were set at 100 and 0% to allow for comparisons between experiments and between neurons. For each treatment and construct tested, FRAP was analyzed for at least 7 neurons across 3 separate transfections.

#### Statistical Analyses

*GraphPad Prism* software package (La Jolla, CA) was used for statistical analyses. All experiments were performed in at least triplicate with statistical tests and post-hoc analyses as indicated in the results section. *P* ≤ 0.05 was considered as statistically significant.

## SUPPLEMENTAL FIGURE LEGENDS

**Figure S1:**
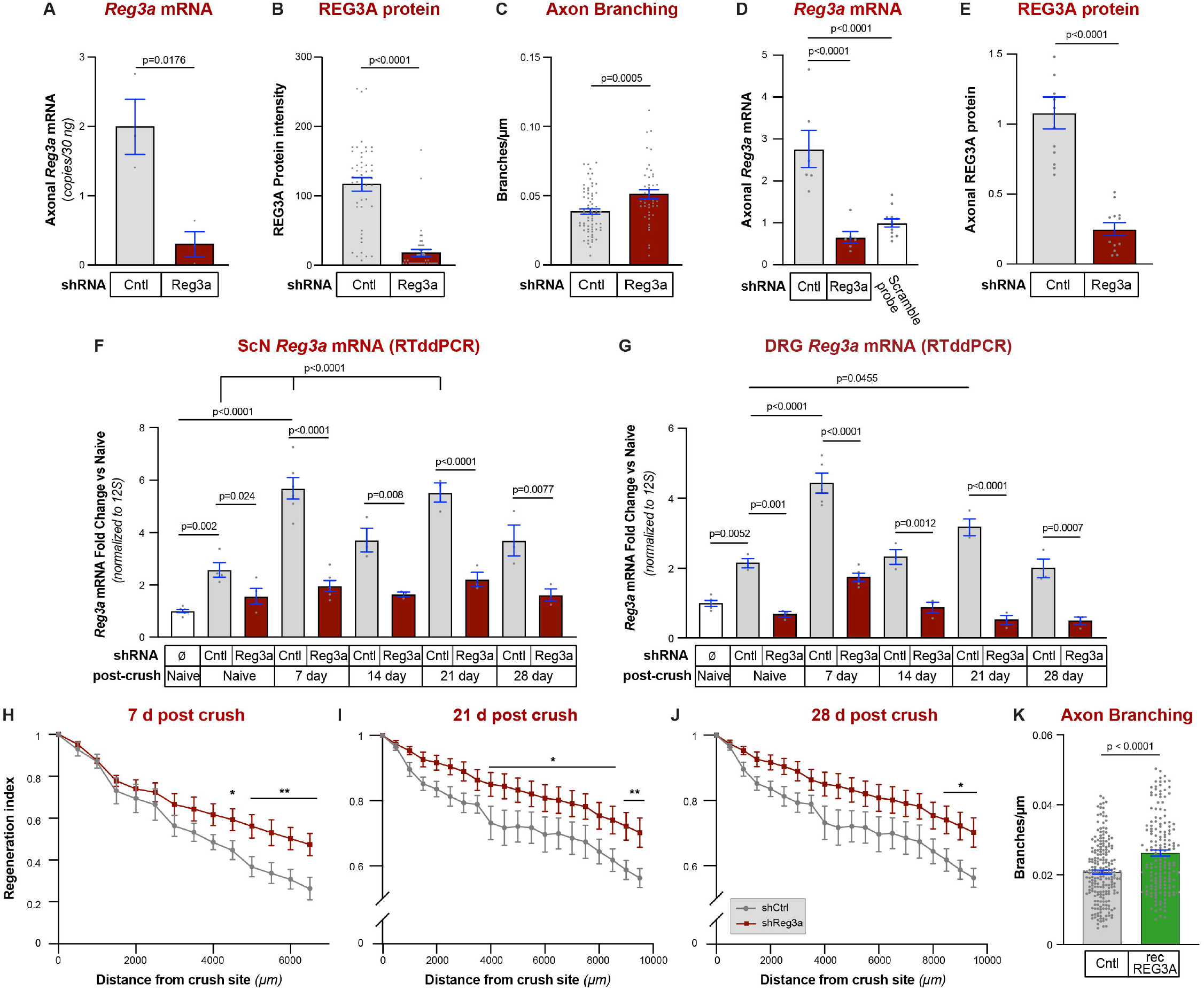
REG3A slows axonal growth. **A-B**, RTddPCR and immunofluorescence data show depletion of *Reg3a* mRNA (**A**) and REG3A protein (**B**) in shReg3a transduced DRG cultures as in Figure 1C-D. Student’s t-test, mean ± SEM (n = 3 mice for A and 45 neurons across ≥ 3 replicate cultures for B). **C**, Quantitation of axon branching in AAV-shCntl vs. -shReg3a transduced DRG cultures from Figure 1C-D shown as mean ± SEM. Student’s t-test (n ≥ 75 neurons across ≥ 3 replicate cultures). **D-E** smFISH and IF data for sciatic nerve sections from mice transduced with AAV-shRNAs show decreased axonal *Reg3a* mRNA (**D**; scramble smFISH probe was used as negative control) and protein (**E**) with shReg3a vs. shCntl. Student’s t-test, mean ± SEM (n = 6 mice per group). **F-G**, RTddPCR for *Reg3a* mRNA in sciatic nerve (ScN) (**F**) and L4-6 DRG lysates (**G**) for naïve, non-transduced (ø) and shReg3a vs. shCntl transduced mice at 0-28 d post-ScN crush corresponding to Figure 1E-J and Suppl. Figure 1H-J. One-way ANOVA with Tukey post-hoc analysis, mean ± SEM (n ≥ 3 per group). **H-J**, Regeneration indices based on SCG10 immunofluorescence for AAV-shRNA transduced mice at indicated post-crush intervals as in Figure 1E-J. Two-way ANOVA with Sidak post-hoc analysis; mean ± SEM; ^*^ p ≤ 0.05 ^**^ p ≤ 0.005 (n ≥ 5 per group). **K**, Quantitation of axon branching in recREG3A vs. Cntl treated DRG cultures from Figure 1L shown as mean ± SEM. Student’s t-test, mean ± SEM (n ≥ 163 neurons across ≥ 3 replicate cultures).

**Figure S2:**
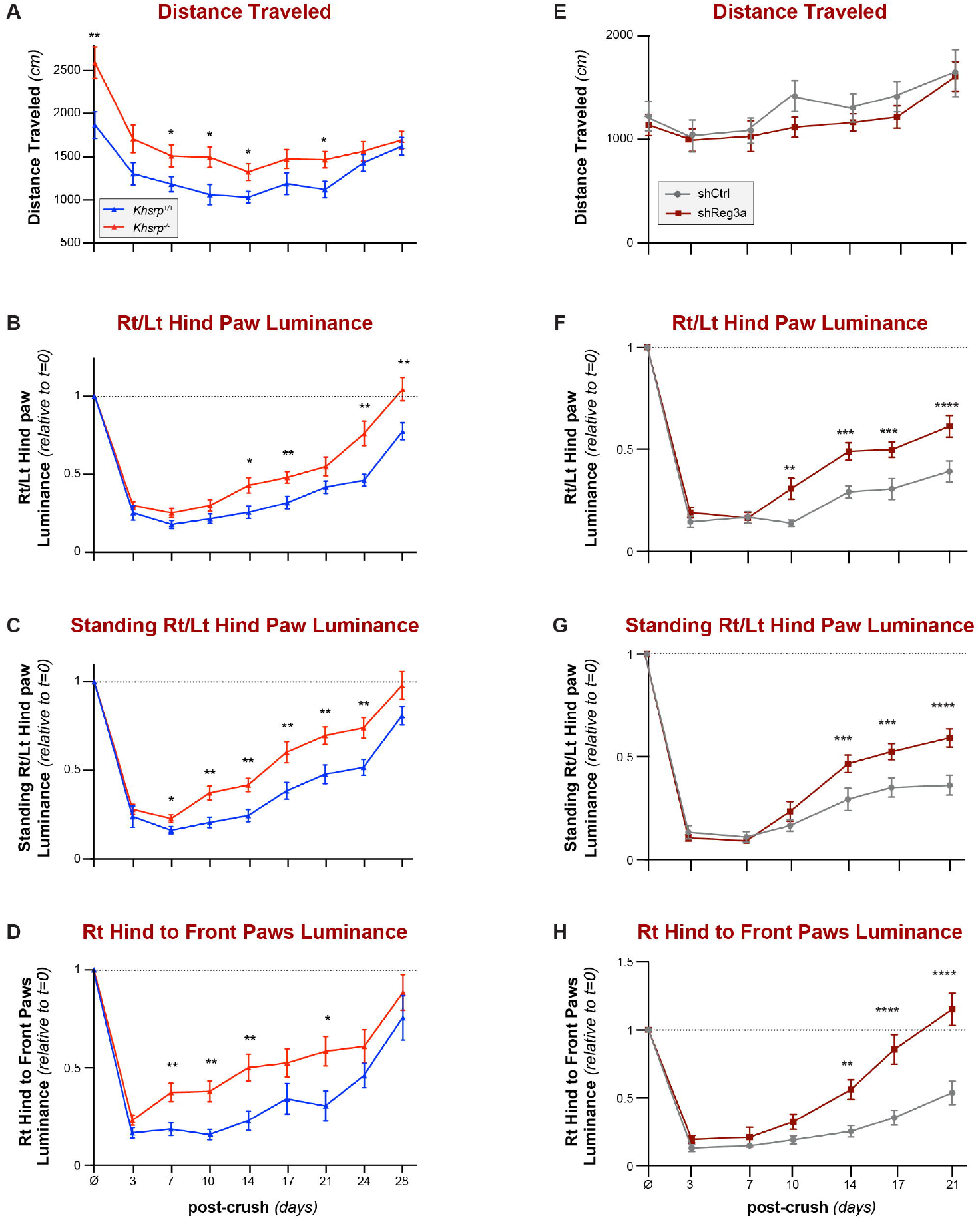
REG3A depletion accelerates functional recovery after nerve crush injury. **A-D**, BlackBox One analyses of *KHSRP*^*-/-*^ compared to *KHSRP*^*+/+*^ mice show increased distance traveled overall (**A**) and, after right-sided ScN crush, increased utilization of the hind paw ipsilateral to ScN crush injury compared to contralateral paw (**B-C**, Rt/Lt hind paw luminance and standing Rt/Lt hind paw luminance, respectively) and average front paws (**D**, Rt hind to front paws luminance). Two-way ANOVA with Tukey post-hoc analysis; mean ± SEM; ^*^ p ≤ 0.05, ^**^ p ≤ 0.005 (n = 18 mice per genotype). **E-H**, BlackBox One analyses for AAV-shCntl vs. -shReg3a transduced mice following unilateral (right) ScN crush injury. There is no difference in distance traveled (**E**), but *Reg3a* depleted mice show increased ipsilateral to injury vs. contralateral hind paws luminance (**F-G**, Rt/Lt hind paw and standing Rt/Lt hind paw luminance), ipsilateral to injury vs. average front paws luminance (**H**) compared to shCntl-transduced mice as in Figure 1G. Two-way ANOVA with Tukey post-hoc analysis; mean ± SEM; ^*^ p ≤ 0.05, ^**^ p ≤ 0.005, ^***^ p ≤ 0.0005, ^****^ p ≤ 0.0001 (n ≥ 10 mice per group).

**Figure S3:**
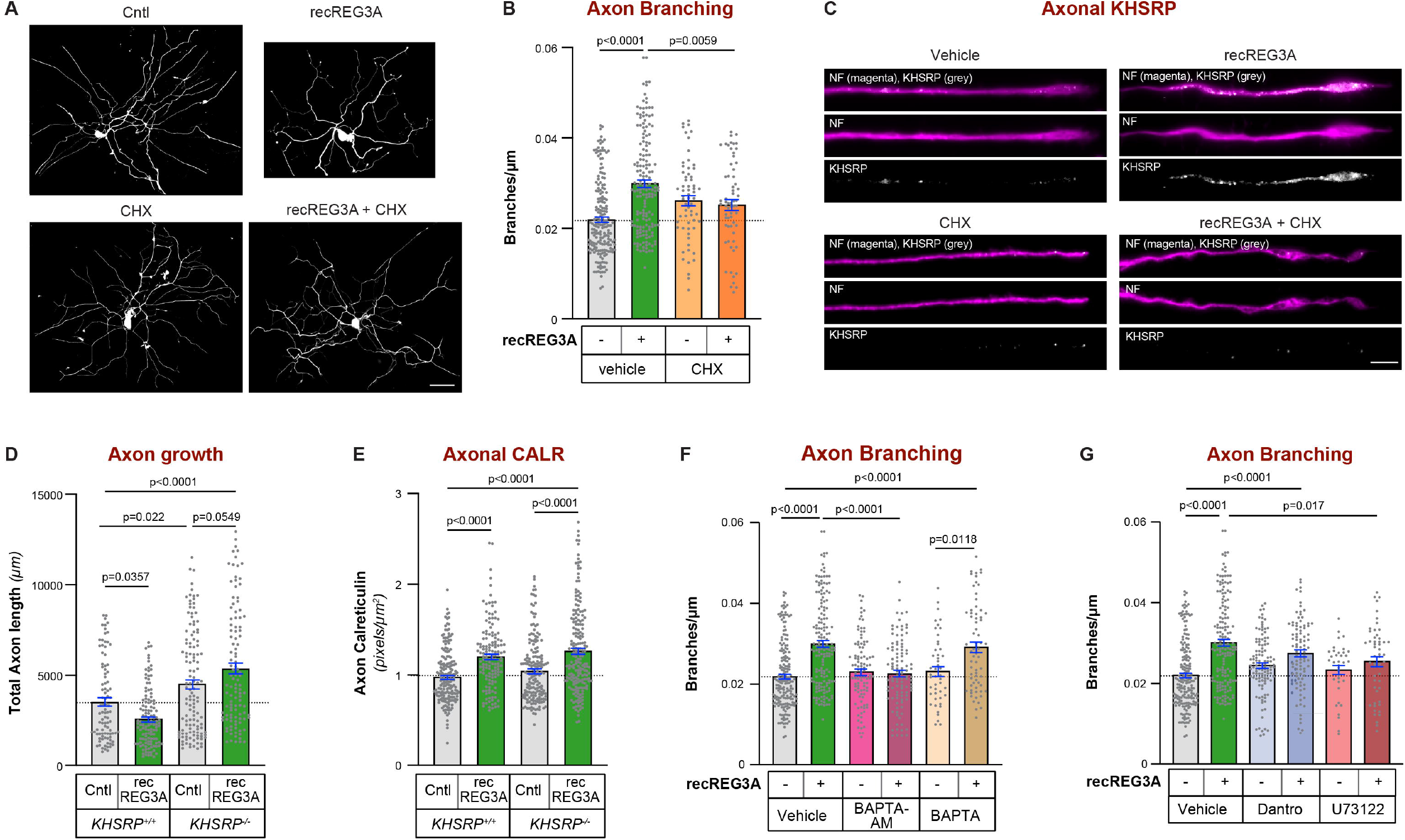
REG3A regulates axon growth and KHSRP levels through modulation of axoplasmic Ca^2^ levels. **A-B**, Representative images for NF + TubJ1 immunofluorescence (**A**) and axon branching quantitation (**B**) for DRGs treated with recREG3A ± cycloheximide (CHX) as in Figure 2A. One-way ANOVA with Tukey post-hoc analysis, mean ± SEM (n ≥ 63 across ≥ 3 replicate cultures); scale bar = 100 µm. **C**, Representative exposure matched images of terminal axons for DRG cultures treated as in A showing SMI312/TuJ1 and KHSRP immunofluorescence corresponding to Figure 2B; scale bar = 5 µm. **D-E**, Axon growth (**D**) and axonal Calreticulin protein (**E**) is shown for DRG neurons cultured from *KHSRP*^*-/-*^ mice ± recREG3A treatment. Neurons lacking KHSRP do not show decrease in axon growth in response to recREG3A but still axonal Calreticulin levels increase in both *KHSRP*^*+/+*^ and *KHSRP*^*-/-*^ DRG neurons. One-way ANOVA with Tukey post-hoc analysis, mean ± SEM (n ≥ 94 neurons in D and n ≥ 143 in E across ≥ 3 replicate cutures). **F-G**, Axon branching in DRG cultures ± recREG3A and BAPTA-AM vs. BAPTA (**F**), or dantrolene vs. U73122 (**G**) corresponding to Figures 3A, and E. One-way ANOVA with Tukey post-hoc analysis, mean ± SEM (n ≥ 46 neurons in F and ≥ 36 in G across ≥ 3 replicate cultures).

**Figure S4:**
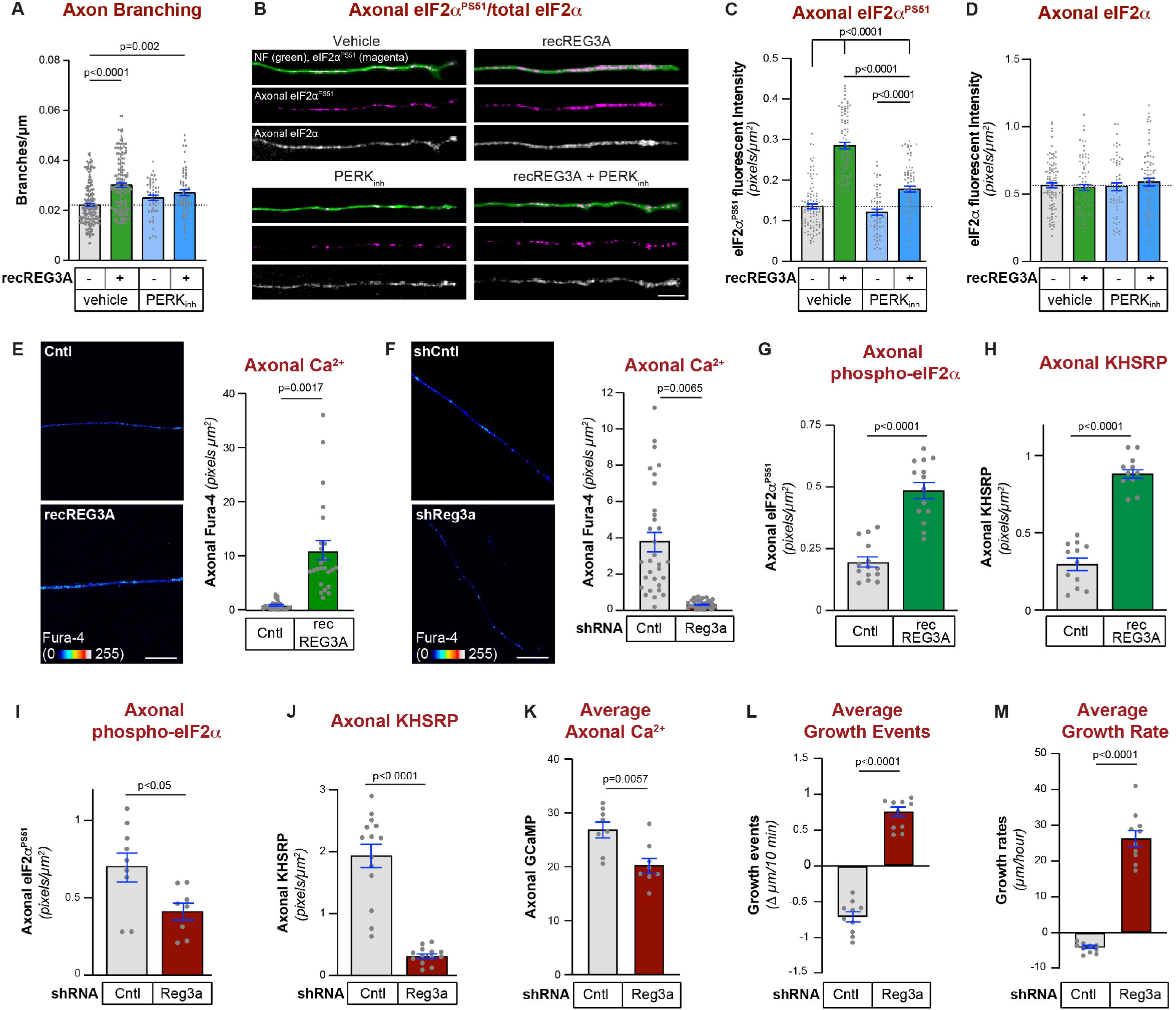
REG3A regulates axon growth and KHSRP levels through modulation of axoplasmic Ca^2^ levels. **A**, Axon branching in DRG cultures ± recREG3A PERK inhibitor corresponding to Figures 3G. One-way ANOVA with Tukey post-hoc analysis, mean ± SEM (n ≥ 60 neurons across ≥ 3 replicate cultures). **B-D**, Representative exposure-matched immunofluorescent images for eIF*α* and eIF*α*^PS51^ (**B**) and eIF*α* and eIF*α*^PS51^ levels in growth cones (**C-D**) of dissociated DRG cultures treated ± recREG3a with PERK inhibitor (PERK_INH_) vs. vehicle (DMSO) corresponding to Figure 3G (n ≥ 60 neurons across ≥ 3 replicate cultures); scale bar = 5 µm. **E-F**, Representative images and quantitations for FURA-4 Ca^2+^-indicator signals in axons of dissociated DRG cultures exposed to recREG3A (**E**) or transduced with AAV-shCntl vs. -shReg3a (**F**). Values shown as mean ± SEM with Student’s t-test (n ≥ 24 neurons over ≥ 3 culture preparations); scale bar = 10 µm. **G-H**, Quantitation of eIF2*α*^PS51^ (**G**) and KHSRP (**H**) immunofluorescence signals in distal axons of intact DRG spot cultures ± recREG3A vs. control for 1 h; mean ± SEM with P values by Student’s t-test (n ≥ 75 neurons over ≥ 3 culture preparations). **I-J**, Quantitation of eIF2*α*^PS51^ (**I**) and KHSRP (**J**) immunofluorescence signals in distal axons of AAV-shCntl vs. -shReg3a transduced DRG spot cultures at 16 h post transection; mean ± SEM with P values by Student’s t-test (n ≥ 75 neurons over ≥ 3 culture preparations). **K**, Average growth cone GCaMPs signal intensity regenerating axons of DRG spot cultures transduced with AAV-shCntl vs. -shReg3a shown as mean ± SEM; P value Student’s t-test (n ≥ 75 neurons over ≥ 3 culture preparations). **L-M**, Average of axon growth events (**L**) and growth rates (**M**) across live cell imaging experiments from Figure 4D-G for regenerating axons in AAV-shCntl vs. -shReg3a transduced DRG spot cultures are shown as mean ± SEM. Student’s t-test, mean ± SEM (n ≥ 75 neurons over ≥ 3 culture preparations).

**Figure S5:**
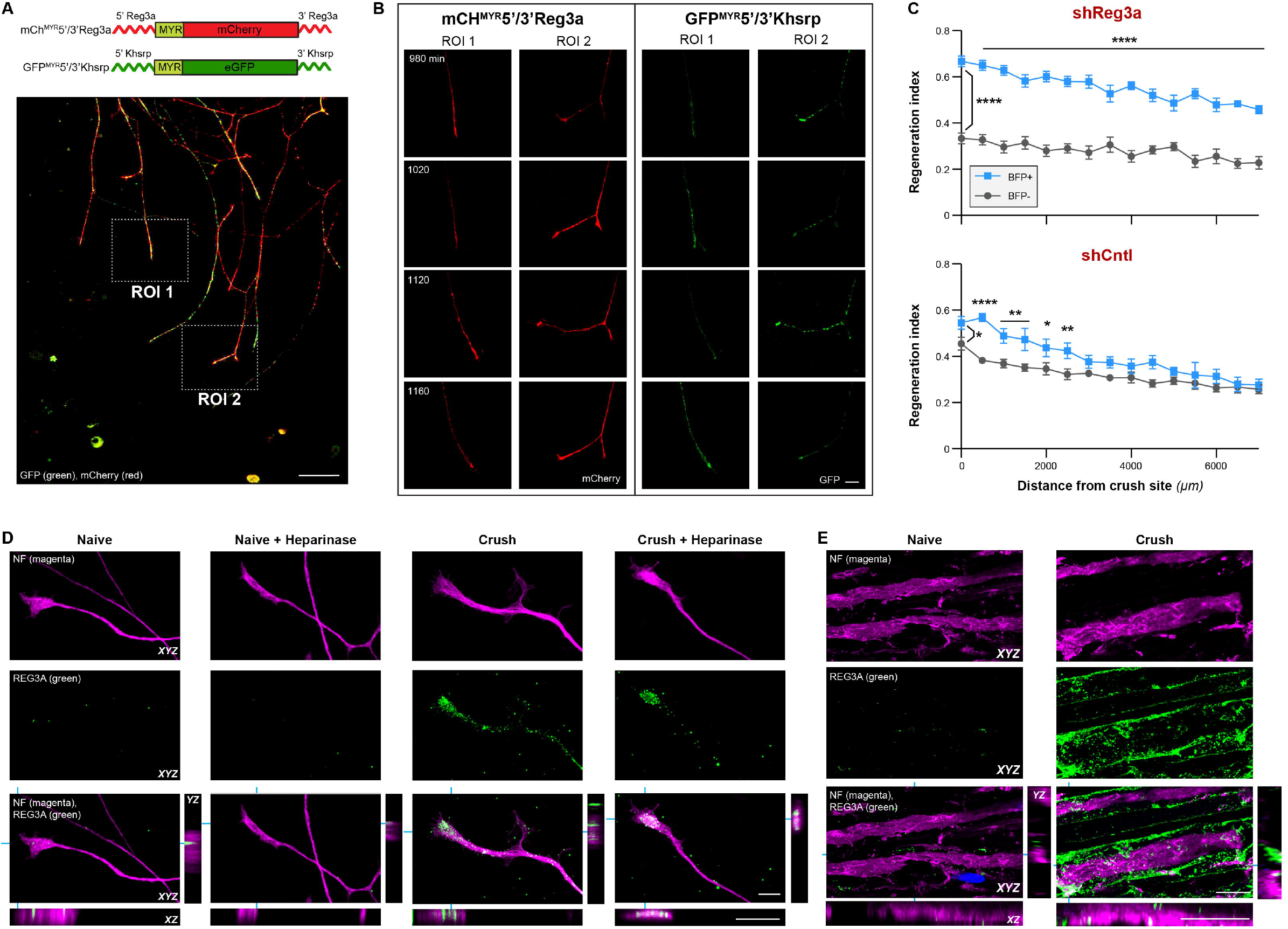
Endogenous REG3A modulates growth cone calcium levels and axon growth events. **A**, Low magnification still image for regions of interest (ROI) from regenerating axons in DRG spot cultures transduced with AAV-GCaMP6s and co-transfected with GFP^MYR^5’/3’Khsrp + mCh^MYR^5’/3’Reg3a that were monitored for distal axon fluorescence in panel B; scale bar = 25 µm. **B**, Representative images for GFP^MYR^5’/3’Khsrp and mCh^MYR^5’/3’Reg3a at indicated time points post-transection are shown for adjacent terminal axons indicated as ROI 1 and 2 in panel A; scale bar = 10 µm. **C**, Regeneration indices of BFP positive vs. BFP negative regenerating axons of shCntl- and shReg3a-transduced mice were identified by SCG10 immunofluorescence from sciatic nerves analyzed in Figure 1E-F. Two-way ANOVA with Sidak post-hoc analysis, mean ± SEM; ^*^ p ≤ 0.05, ^**^ p ≤ 0.005, ^***^ p ≤ 0.0005, and ^****^ p < 0.0001 (n ≥ 5 mice per group). **D**, Representative confocal XYZ maximum projection images for REG3A protein and NF from dissociated L4-6 DRG cultures harvested from naïve vs. 7 d injury-conditioned (‘Crush’) mice. These show increased REG3A protein in the injury-conditioned cultures. REG3A is largely concentrated extracellularly adjacent to the axonal membrane, as emphasized by the inset orthogonal XZ and YZ projections. Heparinase treatment decreases the membrane adjacent REG3A signal, particularly in the injury-conditioned DRG cultures. See Suppl. Video 2 for 3D rendering of the XYZ images shown here; scale bars = 5 µm. **E**, Representative confocal XYZ maximum projection images for optically isolated axons showing REG3A protein and NF in naïve vs. regenerating sciatic nerve (7 d post-crush) with insets showing orthogonal XZ and YZ projections. Note that REG3A is increased in regenerating nerve sections and appears mostly outside of the axon as seen in panel D. See Suppl. Video 3 for 3D rendering of the XYZ images shown here; scale bars = 10 µm.

## SUPPLEMENTAL VIDEOS

**<u>Video S1:</u> *Depletion of Reg3a accelerates axon regeneration***.

Representative video sequences for BFP signals in regenerating axons of shCntl-BFP-vs. shReg3a-BFP-transduced DRG spot cultures as indicated; scale bar = 10 µm.

**<u>Video S2:</u> *Increased REG3A in injury conditioned DRG cultures is largely extracellular***.

Representative 3D renderings of REG3A protein and NF from dissociated L4-6 DRG cultures harvested from naïve vs. injury conditioned (‘Crush’) mice treated with heparinase as indicated; scale bar = 10 µm.

**<u>Video S3:</u> *Increased REG3A in regenerating peripheral nerve is largely extracellular***.

Representative 3D renderings of REG3A protein and NF in naïve vs regenerating sciatic nerve (7 d post-crush); scale bar = 10 µm.

**Table S1:**
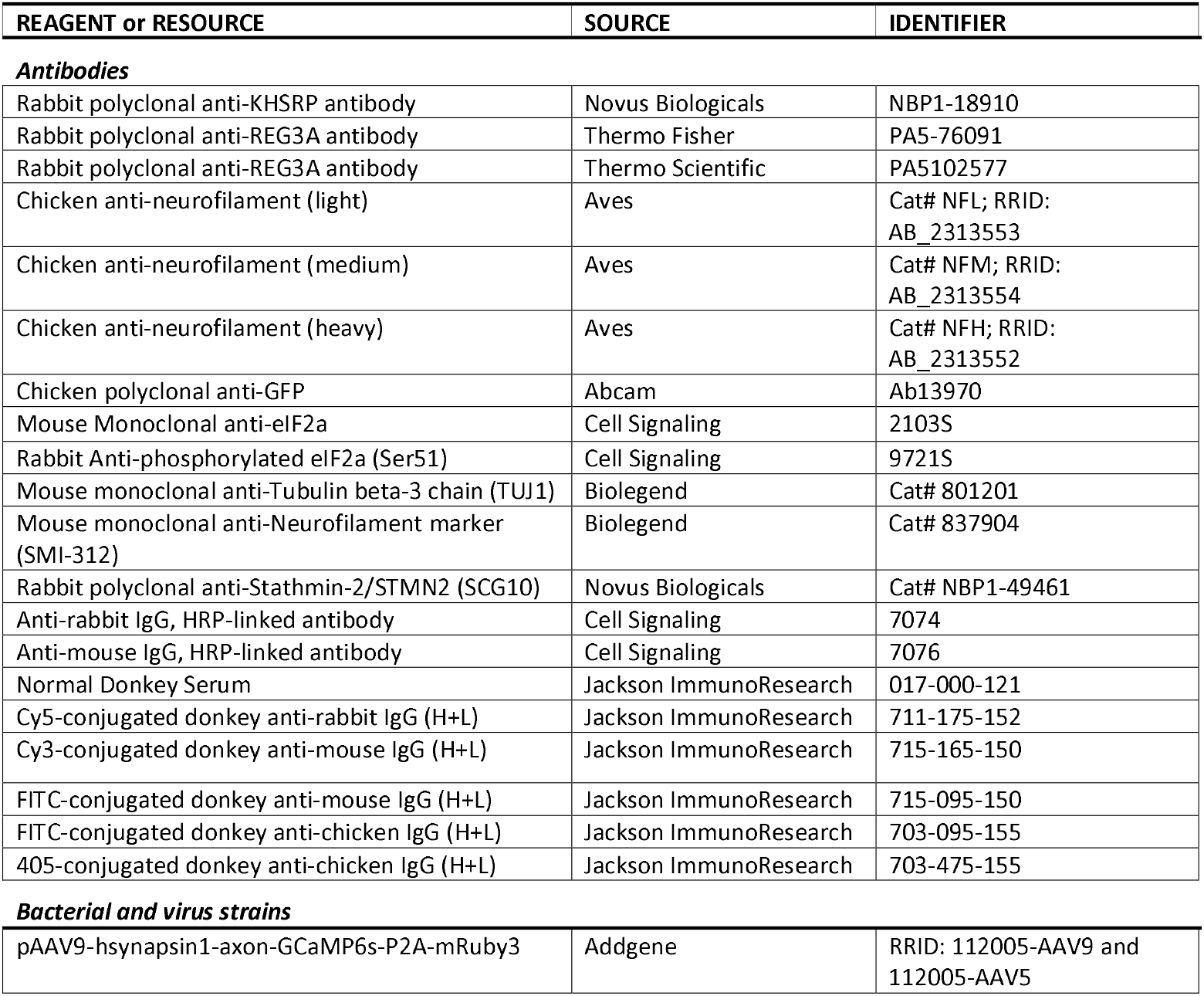
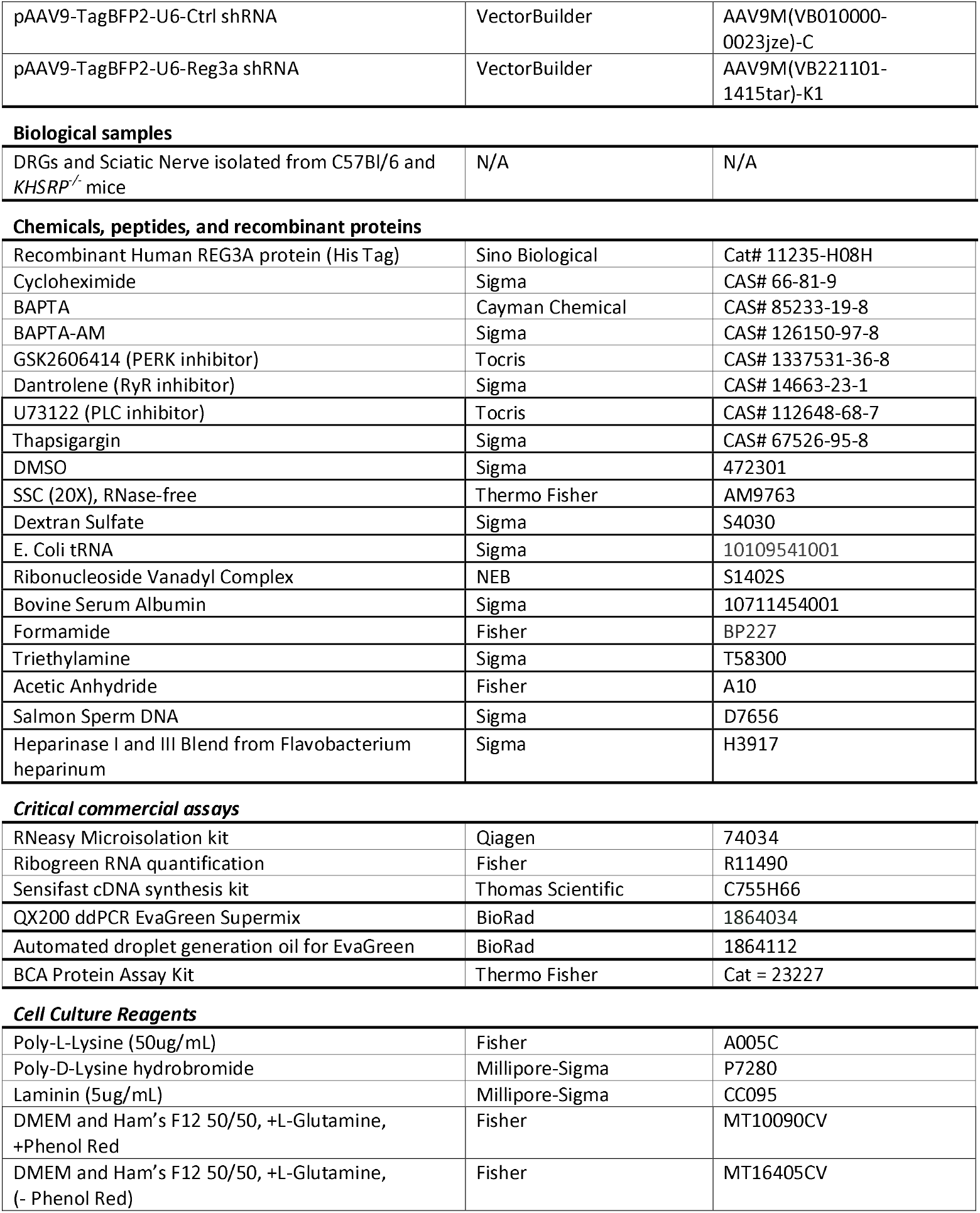
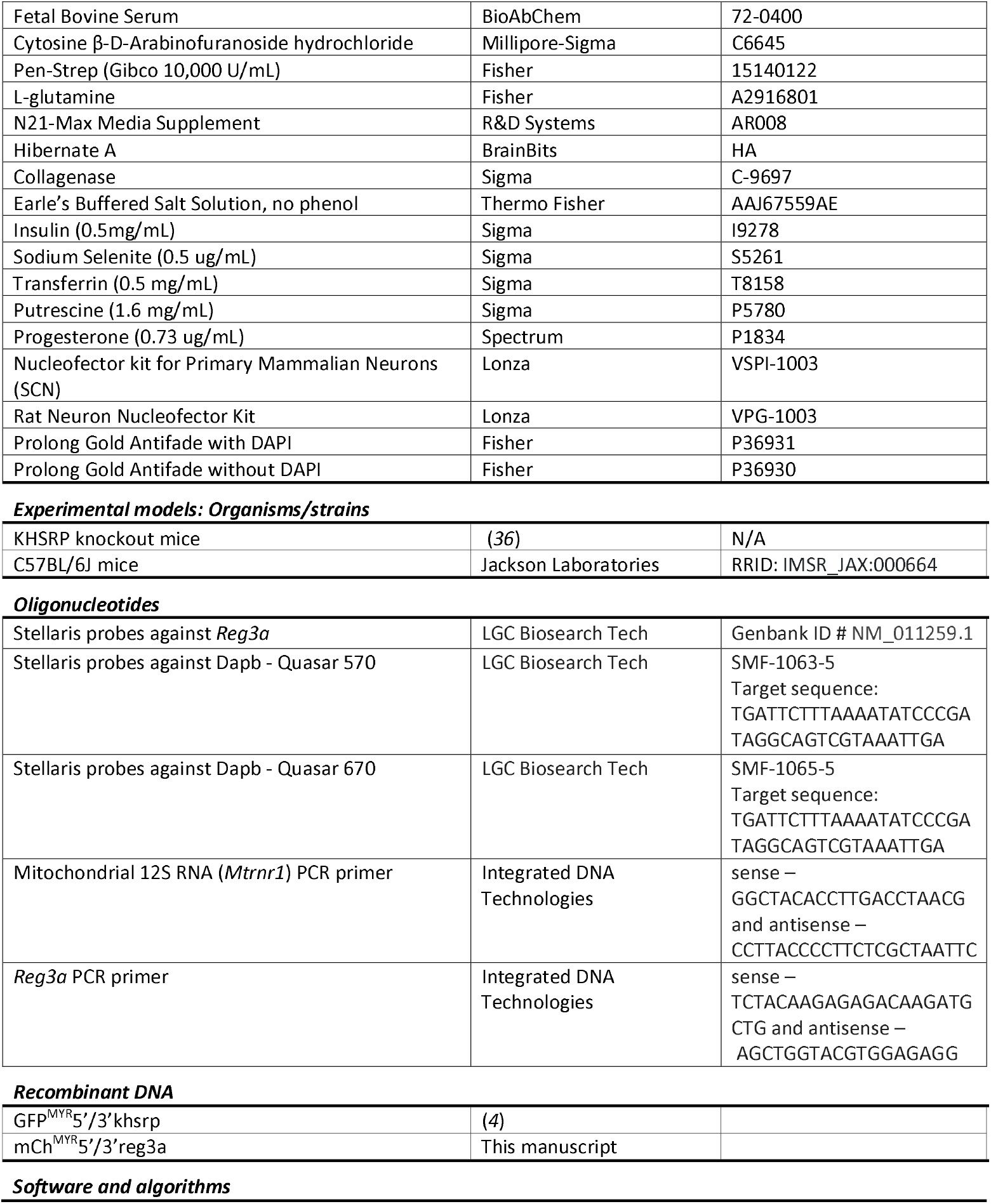
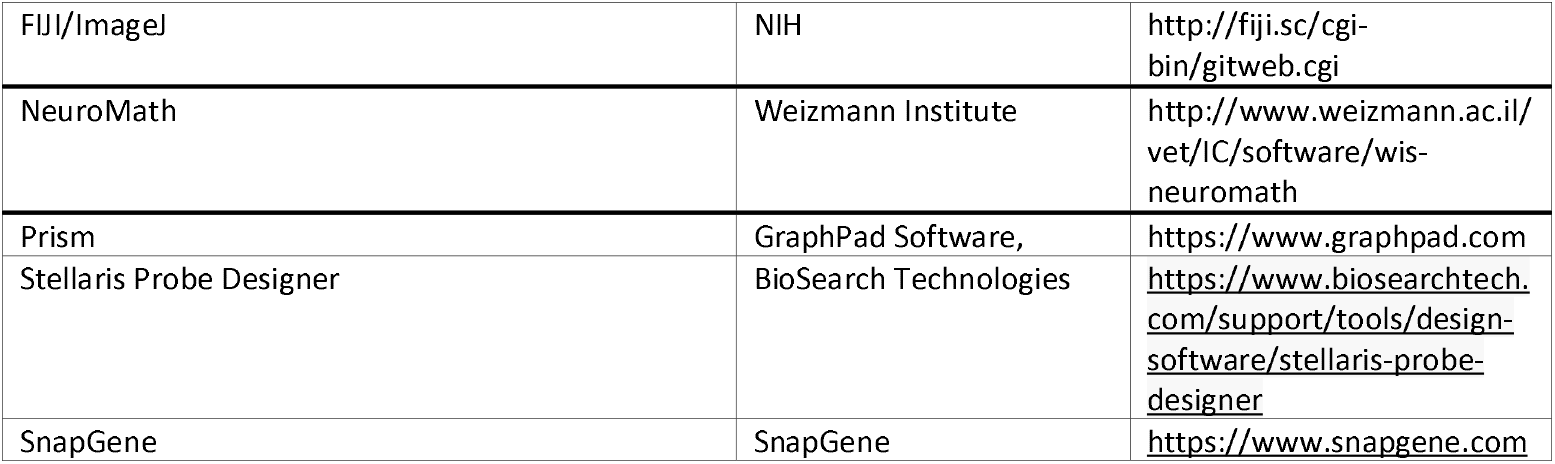
Summary of key reagents and sources.

